# Statistical inference for the evolutionary history of cancer genomes

**DOI:** 10.1101/722033

**Authors:** K. N. Dinh, R. Jaksik, M. Kimmel, A. Lambert, S. Tavaré

## Abstract

Recent years have produced a large amount of work on inference about cancer evolution from mutations identified in cancer samples. Much of the modeling work has been based on classical models of population genetics, generalized to accommodate time-varying cell population size. Reverse-time genealogical views of such models, commonly known as coalescents, have been used to infer aspects of the past of growing populations. Another approach is to use branching processes, the simplest scenario being the linear birth-death process (lbdp), a binary fission Markov age-dependent branching process. A genealogical view of such models is also available. The two approaches lead to similar but not identical results. Inference from evolutionary models of DNA often exploits summary statistics of the sequence data, a common one being the so-called Site Frequency Spectrum (SFS). In a sequencing experiment with a known number of sequences, we can estimate for each site at which a novel somatic mutation has arisen, the number of cells that carry that mutation. These numbers are then grouped into sites which have the same number of copies of the mutant. SFS can be computed from the statistics of mutations in a sample of cells, in which DNA has been sequenced. In this paper, examine how the SFS based on birth-death processes differ from those based on the coalescent model. This may stem from the different sampling mechanisms in the two approaches. However, we also show mathematically and computationally that despite this, they can be made quantitatively comparable at least for the range of parameters typical for tumor cell populations. We also present a model of tumor evolution with selective sweeps, based on coalescence, and demonstrate how it may help in understanding the past history of tumor as well the influence of data pre-processing. We illustrate the theory with applications to several examples of The Cancer Genome Atlas tumors.

## 1 Introduction and preliminaries

The aim of this paper is to present mathematical models that can be used to extract information regarding cancer evolution from the genome sequences of human cancers. This includes the history of growth and mutation and effects such as genetic drift and selective sweeps. Our aim is to point out how mathematical and statistical modeling may help in elucidating problems that frequently have been tackled using intuitive approaches.

Biological cells undergo mutations as they proliferate and such mutations can be neutral, advantageous, or deleterious. The rate of mutation depends on the environment and DNA repair mechanisms. Progress in genome sequencing has allowed cataloguing not only reference genomes of many biological species but also of variants characteristic of human, animal and plant diseases. In particular, initiatives such as the The Cancer Genome Atlas program and the International Cancer Genome Consortium have allowed determination of sets of genomic variants characteristic of some 50 human tumors, with several hundred specimens of each, thus detailing their common mutational features.

One difficulty that arises is that most of the genome sequences available result from so-called bulk sequencing, in which DNA from a sample of cells obtained from the tumor and its environment is cut into fragments, amplified and sequenced, resulting in reads that are aligned with the human reference genome. The resulting genome sequence includes variants that are characteristic of different but not easily identifiable sub-populations of tumor cells. Short of sequencing a representative subset of genomes of individual cells, this difficulty cannot at present be radically improved. Nevertheless, bulk-sequencing data constitute most of the material currently available and it seems important to try to understand the message they carry regarding tumor origin and natural course, perhaps distorted by treatment. This might be called “the genomic archaeology of tumors”.

There are two principal issues arising in the analysis of bulk sequencing data from a tumor: the choice of a model for cell division, and the choice of a model for the way in which the cells are sampled.

Recent years have produced a large amount of work on inference about cancer evolution from mutations identified in cancer samples (cf. Nowell (1976), Greaves and Maley (2012), Sottoriva et al. (2013, 2015), Williams et al. (2018)). Much of the modeling work has been based on classical models of population genetics, generalized to accommodate time-varying cell population size. Reverse-time, genealogical, views of such models, commonly known as coalescent theory, have been used to infer aspects of the past of growing populations. Another approach is to use branching processes, the simplest scenario being the linear birth-death process (lbdp), a binary fission Markov age-independent branching process. A genealogical view of such models is also available. As will be seen in the sequel, the two approaches lead to similar but not identical results.

The “population” in the models we discuss is the collection of all cells in a given tumor. These cells are sampled (for example, through a biopsy) and the DNA they contain is sequenced. Typically a so-called normal DNA sample from the patient is obtained, and a comparison results in somatic variant DNA sites being determined. These variants are based on a sample of reads that is quite difficult to characterize, one reason being that the reads represent a mixture of variants present in different cells of the tumor. We will present some simple models that reflect sampling and show how they work on simulated and real data.

Inference from evolutionary models of DNA often exploits summary statistics of the sequence data, a common one being the so-called Site Frequency Spectrum. In a sequencing experiment with a known number of sequences, we can estimate for each site at which a novel somatic mutation has arisen, the number of cells that carry that mutation. These numbers are then grouped into sites which have the same number of copies of the mutant. Figure 1 gives an example; time is running down the page. The genealogy of a sample of *n* = 20 cells includes 13 mutational events. We can see that mutations 4, 5, 7, 10, 11, 12, and 13 (a total of 7 mutations) are present in a single cell, mutations 1, 2, and 3 (total of 3 mutations) are present in 3 cells, mutations 8 and 9 (a total of 2 mutations) are present in six cells, and mutation 6 is present in 17 cells. If we denote the number of mutations present in *k* cells by *S*_*n*_(*k*), we see that in this example, *S*_*n*_(1) = 7, *S*_*n*_(3) = 3, *S*_*n*_(6) = 2, and *S*_*n*_(17) = 1, with all other *S*_*n*_(*j*) equal to 0. The vector (*S*_*n*_(1), *S*_*n*_(2), …, *S*_*n*_(*n* − 1)) is called the (observed) Site Frequency Spectrum, abbreviated to SFS. It is conventional to include only sites that are segregating in the sample, that is, those for which the mutant type and the ancestral type are both present in the sample at that site. Mutations that occur prior to the most recent common ancestor of the sampled cells will be present in all cells in the sample; these are not segregating and are called truncal mutations.

**Figure 1:**
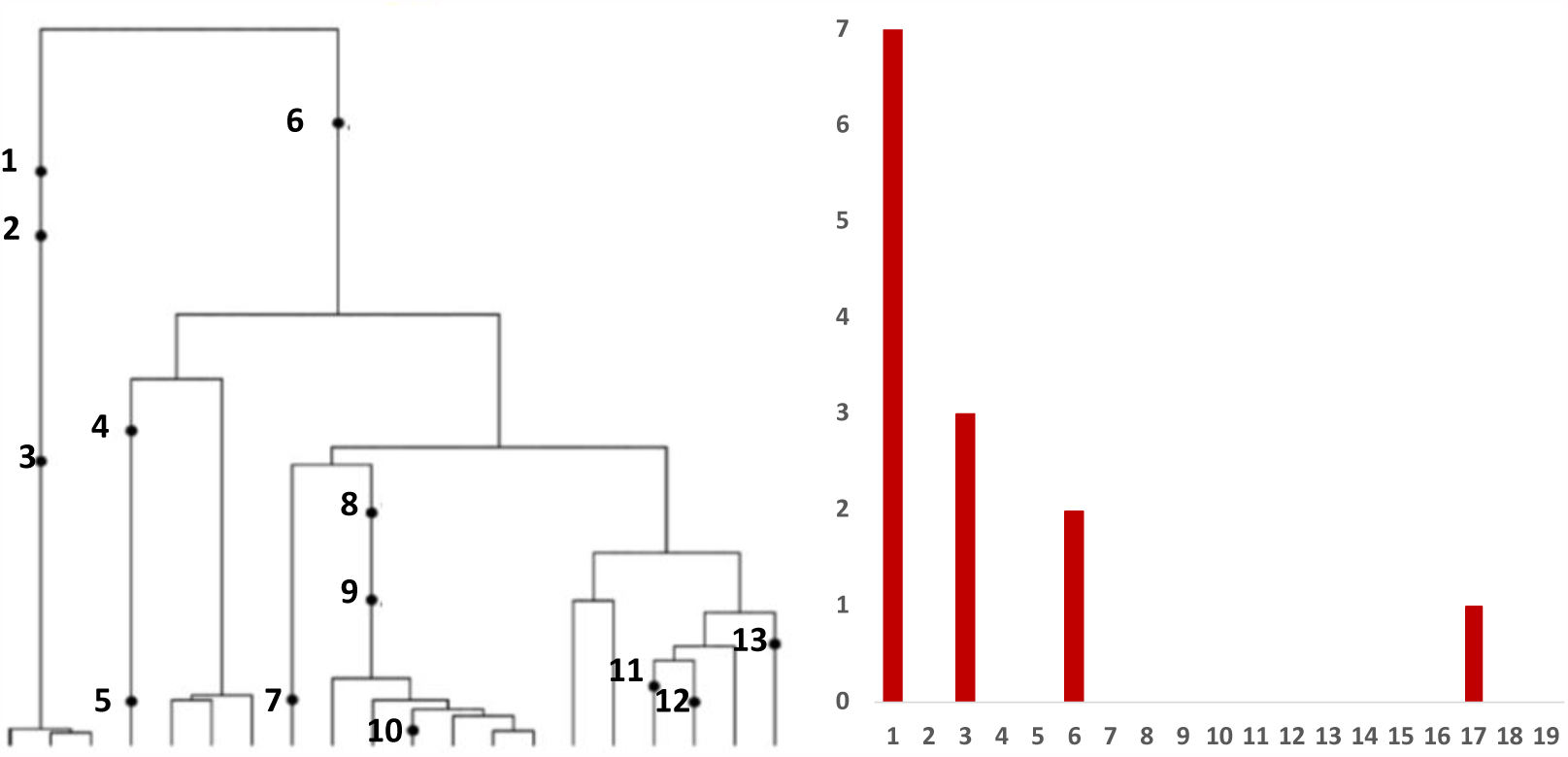
Left panel: Genealogy of a sample of *n* = 20 cells includes 13 mutational events, denoted by black dots. Mutations 4, 5, 7, 10, 11, 12, and 13 (total of 7 mutations) are present in a single cell, mutations 1, 2, and 3 (total of 3 mutations) are present in three cells, mutations 8 and 9 (2 mutations) are present in six cells, and mutation 6 (1 mutation) is present in 17 cells. Right panel: The observed site frequency spectrum, *S*_20_(1) = 7, *S*_20_(3) = 3, *S*_20_(6) = 2, and *S*_20_(17) = 1, other *S*_*n*_(*k*) equal to 0.

In most cancer sequencing experiments, we do not know the number of sequences that were sampled. Nonetheless, we can estimate the relative proportion of the mutant at each segregating site, and so arrive at a frequency spectrum based on proportions. We continue to use the term SFS for such a spectrum, as there should be no cause for confusion.

The emphasis in the definition of the SFS is that it is based on a DNA sample extracted from cells, which does not usually constitute the entire tumor population. Moreover, at any DNA site, the sample can, and most frequently does, arise from DNA of different cells, as will be explained in Section 5. This underscores the importance of developing a sampling theory for the SFS estimated from genome sequencing data. We will develop some simple results in Section 6.

## 2 Modeling exponentially growing cell populations

Stochastic models of growth and inheritance in biological populations follow two major traditions, one originating from population genetics, the other from population dynamics. Population genetics models, including models of Wright and Fisher, Moran, and Cannings, assume in their original form time-constancy of the population size. Under this assumption, major mathematical population genetics results such as the Ewens Sampling Formula (Ewens, 1972), Kingman’s coalescent (1982a, 1982b), Kimura’s use of diffusion approximations (reviewed in Watterson (1996)) and many others, have been derived. The tradition from which the constancy assumption stems underscores the importance of constraints under which populations evolve, such as space and resource limitations for populations of animals and human, or hormonal controls and tissue size bounds for cell populations in multi-cellular organisms.

The population dynamics tradition, embodied by branching process models, emphasizes growth and stochastic fluctuations stemming from birth and death events of a finite collection of independent individuals (here, cells). Historically, models such as these have been employed to reproduce growth of bacterial populations or other cells in culture, the growth of cancerous tumors, or to develop methods for estimation of mutation rates, such as Luria and Delbrück’s fluctuation analysis (Luria and Delbrück (1943), Lea and Coulson (1949)); see also Gerrish (2008).)

How can we align these two rather different approaches? One way is to relax the constancy assumption in the population genetics models, and this will be the first type of model discussed in this section. If the population size is growing exponentially in time, this model can be compared to the supercritical branching process (cf. Jagers (1975), and Haccou et al. (2005)). There are three differences remaining: first, the supercritical branching process grows exponentially only in the limit (and in expectation); second, the “population growth rate” of the coalescent is a summary parameter that may correspond to a wide range of supercritical branching models with different population size distributions; and third, in birth-death processes, coalescent events coincide with population size increments. It is therefore of interest to know how these two methods compare when applied to simulated or real cell populations. Another, potentially major, difference is that in the coalescent models we assume we can trace the sampled cells back to their most recent common ancestor. There are several different sampling versions for the branching process. The difference will become transparent later on.

We will compare in this section two models based on the genealogical view of cell evolution, the first one being the variable population size coalescent. The other is an analogous reverse-time, genealogical approach, known as the coalescent point process, which is based on the linear birth and death process, mathematically equivalent with the branching process with binary fission and exponentially distributed cell lifetimes; cf Kimmel and Axelrod (2015).

We first describe both approaches in general terms, and then compare the expressions for site frequency spectra (SFS) under these approaches.

### 2.1 A Moran model for cell division

The simplest model for cell division in a constant-size population of *N* cells is the Moran model (Moran (1958, 1962)). We describe the process backwards in time, noting that there are several essentially equivalent methods for doing this. Such a description is convenient for simulating the effects of mutation on the cells in the sample, and leads to the study of the ancestral process that counts the number of distinct ancestral cells in the history of the sample back to its MRCA. Imagine, then, that birth-death events occur independently to cells at rate 1. At one of these events, one cell dies, and another is chosen from the remaining *N* − 1 to divide. If there are currently *i* distinct ancestral cells in a sample of size *n*, then the next event in the past results in *i* − 1 distinct ancestors if, and only if, the pair of individuals is in the sample of *i*, an event of probability 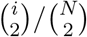. Thus the rate at which the number of distinct ancestors reduces by 1 is

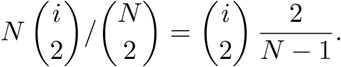

It is convenient to consider what happens for large populations of cells. If time is scaled in units of *N/*2, then asymptotically as *N* → ∞, the ancestral process drops from *i* to *i* − 1 at rate 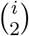, resulting in a particularly simple ancestral process known as the *coalescent*.

To describe the ancestral process, let *T*_*n*_, *T*_*n*−1_, …, *T*_2_ denote the lengths of time during which the sample has *n, n* − 1, …, 2 distinct ancestors back in time to its most recent common ancestor. Kingman (1982b) showed that the *T*_*j*_ are independent exponential random variables, with

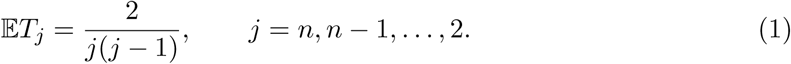

The Markov chain {*A*_*n*_(*t*), *t* ≥ 0} that counts the number of distinct ancestors of the sample a time *t* ago has transition rates

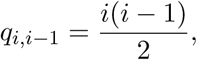

staying at 1 when the sample has been traced back to its most recent common ancestor.

The variable population size version of this model supposes that at time *t* ago, the population size is *N λ*_*N*_(*t*). Arguing as above, the rate at which *i* ancestral lines coalesce to *i* − 1 is

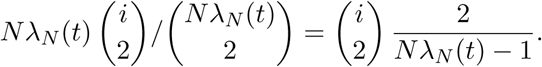

Scaling time in units of *N/*2, as in the constant size case, we see that in the limit as *N* → ∞, the rate at time *s* becomes 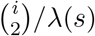, where *λ*(*s*):= lim *λ*_*N*_(*N s/*2).

In the setting of exponential growth from the past, we have *λ*_*N*_(*t*) = exp(*−rt*), so that

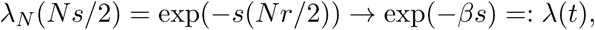

where we have assumed that *N r/*2 *→ β* as *N* → ∞. This process maintains the random merging of ancestral lines back into the past, but the distribution of the coalescence times 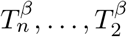 is more complicated, and most easily described by the fact that the ancestral process 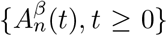 for the exponential model results from a deterministic time change of the constant size case:

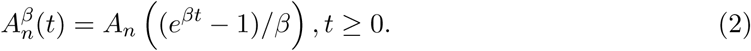

We use this fact to simulate the 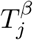, as shown in Section A.1.

### 2.2 A branching process model for cell division

Lambert (2010) and Lambert and Stadler (2103) demonstrated that under general assumptions on birth and death rates, the coalescent of a binary branching process has iid coalescence times. More specifically, if the branching process was started from one cell at time 0 and conditioned to have at least one cell alive at time *t*, then the coalescent tree of the *N*_*t*_ ≠ 0 cells alive at *t* is a coalescent point process (abbreviated CPP): that is, the *N*_*t*_ − 1 coalescence times form a sequence of independent copies of some rv *H* whose law can be characterized in terms of the birth and death rates of the process, killed at its first value larger than *t*. This conclusion hinges on the manner the tree is ordered, the two rules being that (1) progeny branch out on the right of the parent, and (2) given progeny’s life-line is on the right of all further descendant of the parent cell (as in the example in Fig. 13). It is common to characterize *H* through its so-called inverse tail distribution *W*,

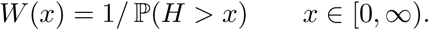

The most general assumptions under which the last statement holds are (i) the per-cell birth (division) rate depends only on absolute time, and (ii) the per-cell death rate depends only on absolute time and cell age (or any other non-heritable trait).

Implicit in condition (2) is that upon division, one can distinguish between the mother cell (whose age continues to increase after division) and the daughter cell (whose age is 0 at division). Another way of expressing this is that cells can have lifetimes that follow a general distribution (not necessarily exponential, possibly even deterministic) which possibly depends on their absolute birth time.

A consequence of the CPP representation is that *N*_*t*_ follows a geometric distribution with failure probability 1/*W*(*t*). Then conditional on *N*_*t*_ = *n*, the coalescence times are iid rvs distributed as *H* conditioned on *H < t*.

A useful feature of CPPs is that a Bernoulli sample from a CPP is again a CPP. More specifically, if each tip of a CPP tree with inverse tail distribution *W* is sampled independently with probability *p*, the tree spanned by the sampled tips is a CPP with inverse tail distribution *W*_*p*_ given by

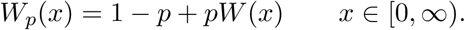

In the case when the bp has constant birth rate *b* and death rate *d* (linear birth-death process), growth rate *r*:= *b − d*,

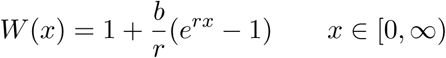

if *b* ≠ *d* and *W* (*x*) = 1 + *bx* if *b* = *d*. Note that in the subcritical case when *b < d*, *P* (*H* = +∞) = 1 *− b/d*.

The coalescent point process representation of reconstructed trees generated by an lbdp is originally due to Popovic (2004) in the critical case (*b* = *d*) and has been extended to non-critical cases and to some non-Markovian branching processes by Lambert (2010) and Lambert and Stadler (2013). A corollary of this representation is that conditional on the number of tips, branching times are independent with an explicit distribution. Note that this corollary is already present in Thompson (1975), Nee et al (1994), Rannala (1997) and Gernhard (2008).

## 3 Site frequency spectra under the infinitely-many-sites model

We examine how the SFS based on birth-death processes differ from those based on the coalescent model. This may stem from the different sampling mechanisms in the two approaches. However, we also show that despite this, they can be made quantitatively comparable at least for the range of parameters typical for tumor cell populations.

### 3.1 The SFS for the coalescent

We assume an infinitely-many-sites (IMS) model of mutation: think of the DNA sequence as a unit interval, and label mutations by a sequence of independent uniform(0,1) random variables. Mutations are almost surely distinct, giving rise to the term “infinitely-many-sites”.

We assume that mutations arise over the lifetime of a cell according to a Poisson process of rate *τ*_*N*_, conditional on the lifetime. The expected number of mutations accumulated per unit time is therefore *τ*_*N*_. If time is rescaled in units of *N/*2, the expected number becomes *N τ*_*N*_/2, so to balance mutation and drift we assume that *N τ*_*N*_ → *ϑ* as the population size increases. To summarize, time is measured in units of *N/*2, mutations occur according to a Poisson process of rate *ϑ/*2 during a cell’s lifetime, *N* is assumed very large, and

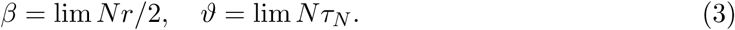

In the large population size limit, mutations take place according to independent Poisson processes of rate *ϑ/*2 on the branches of the coalescent tree, conditional on the lengths of the branches.

Griffiths and Tavaré (1998) provide a general coalescent framework for the expected number 𝔼*S*_*n*_(*k*) of mutant sites having *k* copies of the mutant in a sample of size *n*, drawn from a population with size changing deterministically in the past. We provide a brief account of their results for the case of exponential population growth and describe a useful approximation due to Durrett (2013).

Griffiths and Tavaré showed that

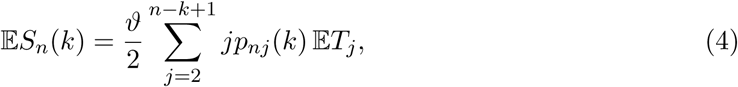

where

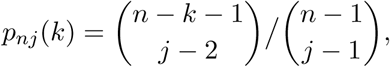

the *T*_*j*_ denoting the coalescence times for the model with exponential growth. While the expectations can be simulated, it is convenient to consider the approximations provided by Durrett (2013), who showed that

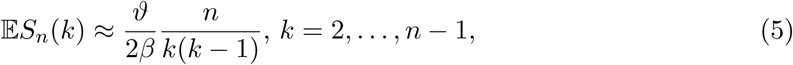

while

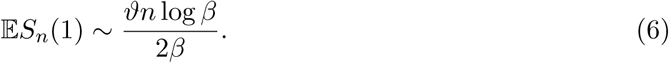

### 3.2 The SFS for the birth-death process

Here, we consider an application of the theory of coalescent point processes to a supercritical linear birth-death process (lbdp) with an ISM mutation model. We derive an explicit expression for the expectation of the site frequency spectrum (SFS) in this case, and develop a simple and efficient simulation scheme based on the CPP representation. In the spirit of the original work, we use unscaled parameters *r* and *θ* here, instead of the scaled *β* and *ϑ*. See Table 2, which displays the conversions.

Lambert (2009) showed that the expected SFS for a sample size *n* has the form

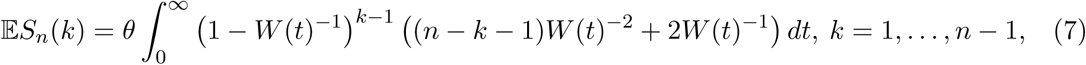

where for the lbdp case the function *W* (*x*) has the form

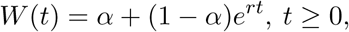

with *r >* 0, *α* ∈ (0, 1), and where *θ* is the mutation rate (the intensity of the Poisson process of mutations assumed in the ISM). Recall that *r* = *b − d* and *α* = 1 *− pb/r*, where *b* is division rate, *d* is death rate and *p* is the fraction of cells sampled. The case when *α* = 0 corresponds to *d* = *b*(1 *− p*), which occurs in particular when *d* = 0 and *p* = 1.

This version of the process is equivalent to the ancestor being “born in the very remote past”, and it is the limit version of the process we will simulate. As will be seen, the discrepancies are small for the cases in which we are interested.

We show in Section A.3 that

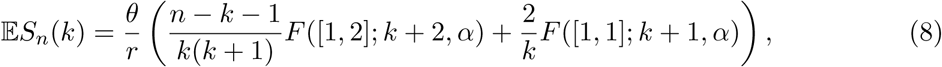

where 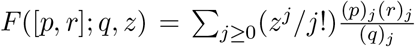 belongs to the hypergeometric family of functions (cf. Abramowitz and Stegun, 1964) and

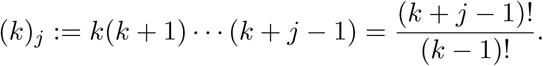

An algorithm for simulating the SFS based on the CPP process is given in Section A.2.

#### 3.2.1 Computational example

We carried out a number of simulation experiments including a range of parameters. Figure 2 depicts results of one such experiment. As can be seen, the average of 10,000 simulated SFS coincides closely with the hypergeometric formula. However, the simulated SFS median becomes equal to 0 for relatively small *k*. For individual SFS, this corresponds to more than SFS half terms being equal to 0, which is consistent with spectra observed in cancer mutations.

**Figure 2:**
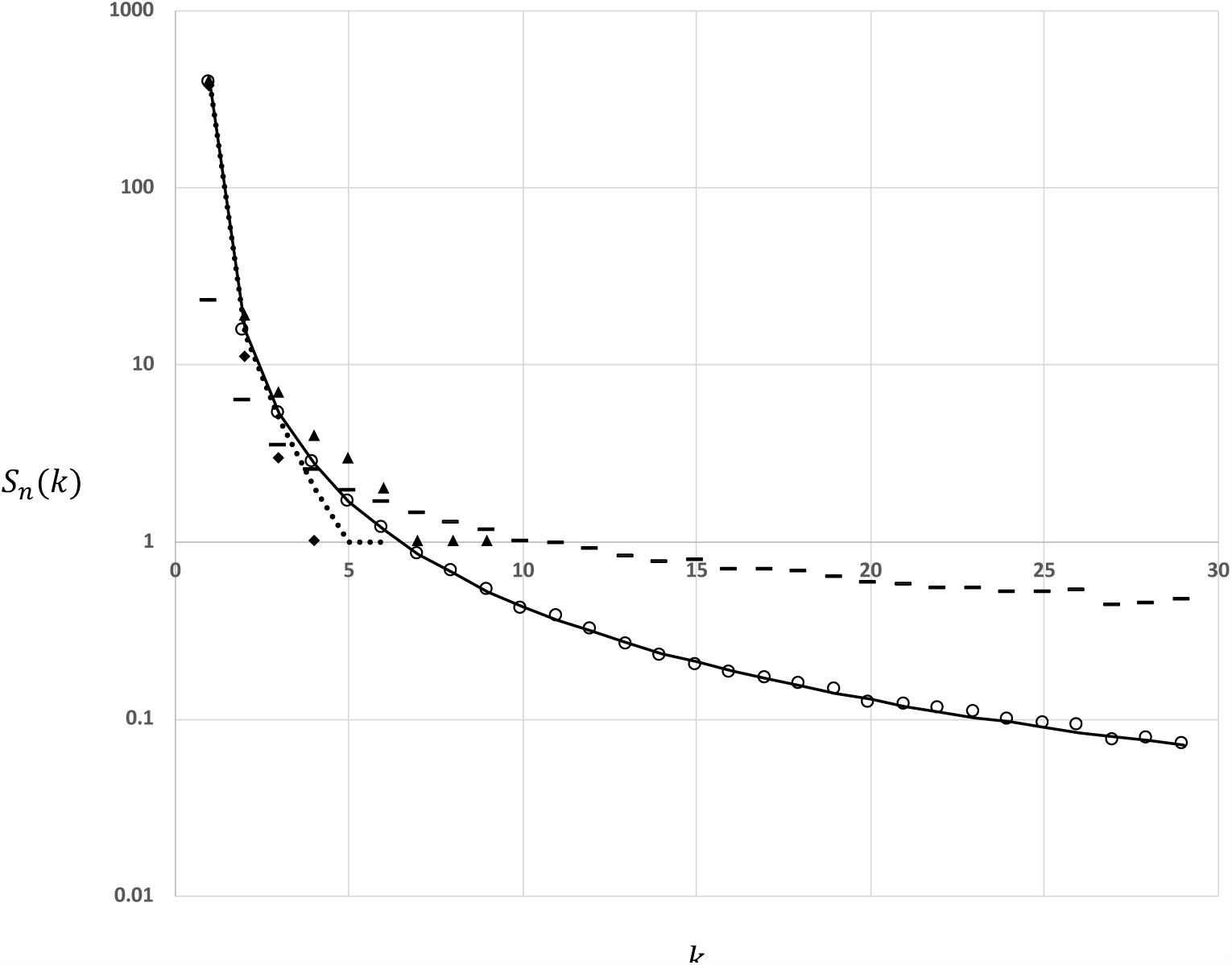
Numerical example of the expected SFS for the lbdp (semi-logarithmic scale). Continuous line: expected SFS 𝔼*S*_*n*_(*k*) (interpolated for visual convenience); circles: corresponding average of 10,000 simulations; dashes: standard deviation estimate based on 10,000 simulations; dotted line, diamonds and triangles: median and first and third quartile of 10,000 simulations. The parameters for this simulation (cf. Table 2) are *n* = *p* 𝔼*N* (*t*) = 30, *r* = 1, *θ* = 1, *α* = 0.999999, *t* = 100 for simulations, *t* = ∞ for 𝔼*S*_*n*_(*k*). Other parameters can be calculated from these.

### 3.3 Using the two coalescents to model tumor growth and mutation

Section 2.1 explains how to use the Moran model with exponentially growing population size to introduce coalescent structure into our cell proliferation model. In the birth-death process approach, we model a growing population of tumor stem cells as a birth-death process in continuous time with parameters *b* and *d*. In biological terms this means that a cell population starts with a single cell at time *t* = 0, the lifetimes of cells are exponentially distributed with parameter *b* + *d*, and that cell divides into two progeny with probability *b/*(*b* + *d*) (probability of self-renewal), or dies with probability *d/*(*b* + *d*). Under these assumptions, the expected cell count at time *t* is equal to 𝔼*N* (*t*) = *e*^*rt*^, *t* ≥ 0, with growth rate *r* = *b − d*.

At time *t* = *x*, when the tumor is diagnosed, its nuclear DNA is sequenced with average coverage *n*. (A more realistic sampling theory appears in Section 6.) This can be represented as binomial sub-sampling from about *N* (*x*) cells with sub-sampling probability *p* = *n/N* (*x*). Notice that the *d*-parameter does not have to be literally equal to the death rate. The model applies equally well to the population of cancer cells, in which case *d* is the combined death and differentiation rate. Mutations occur according to the ISM model, at rate *θ*.

For illustrative purposes, this growth model will be parameterized to reflect several scenarios differing with respect to growth rate and efficiency of division. In the current computations we assume that the tumor is detected when it contains approximately *N* (*x*) = 10^7^ cells. How can we relate it to sizes of human tumors? An analysis of this issue has been published by Del Monte (2009), who addresses the commonly held view that 1cm^3^ tumor contains 10^9^ cells. The author concludes that this is true for “normal” human cell sizes, while tumor cells may frequently be larger and interspersed with other cells, so it may be more appropriate to claim that 1cm^3^ contains 10^8^ or even fewer cells (so that 10^10^ cells might occupy a cube 4.64 cm each side or larger). Ling et al. (2015) consider a 1mm thick slice of hepatocellular carcinoma, roughly a disc 3.5cm in diameter (volume of a cube 0.98cm each side) and apparently assume (see their Table 1) only 10^5^ cells (aside from this, they sample mutations in different tumor regions and find the resulting SMS in agreement with Durrett’s formula, based on non-singletons, which does not relate to *N*). To sum up, our assumed *N* (*x*) = 10^7^ cells seems on target.

**Table 1:** Calculations of parameters for the tumor birth-and-death process

| $x$ | 40 | 400 | 4000 |
| --- | --- | --- | --- |
| $r = \ln[\mathbb{E} N(x)]/x$ | 0.40295 | 0.04029 | 0.00403 |
| $d = b - r$ | 0.59704 | 0.95970 | 0.99597 |
| $1 - \alpha = bp/r$ | $7.44 \times 10^{-6}$ | $7.44 \times 10^{-5}$ | $7.44 \times 10^{-4}$ |

We will consider slow-, moderate-, and fast growing tumors that reach this size within *x* = 4000, 400, and 40 days, respectively. Also, we will assume the surviving cell average lifetime 1*/b* corresponding to *b* = 1 day, which is consistent with the average cell cycle time in mammalian cells (Mura et al., 2018). Calculations of other parameters corresponding to these input specifications are listed in Table 1.

Figure 3 depicts the expected SFS based on the hypergeometric formula (20) with parameters as described above. The three cases considered are depicted along with the corresponding SFS resulting from the Griffiths-Tavaré theory, this latter using scaled parameters as at the top of the present section. In addition, Figure 4 depicts the expected SFS based on the hypergeometric formula (20) with parameters as for the center scenario in Table 1, but with parameter 1 *− α* varying from 10^*−*8^ through 0.5. For comparison, Durrett’s approximation (5) is included.

**Figure 3:**
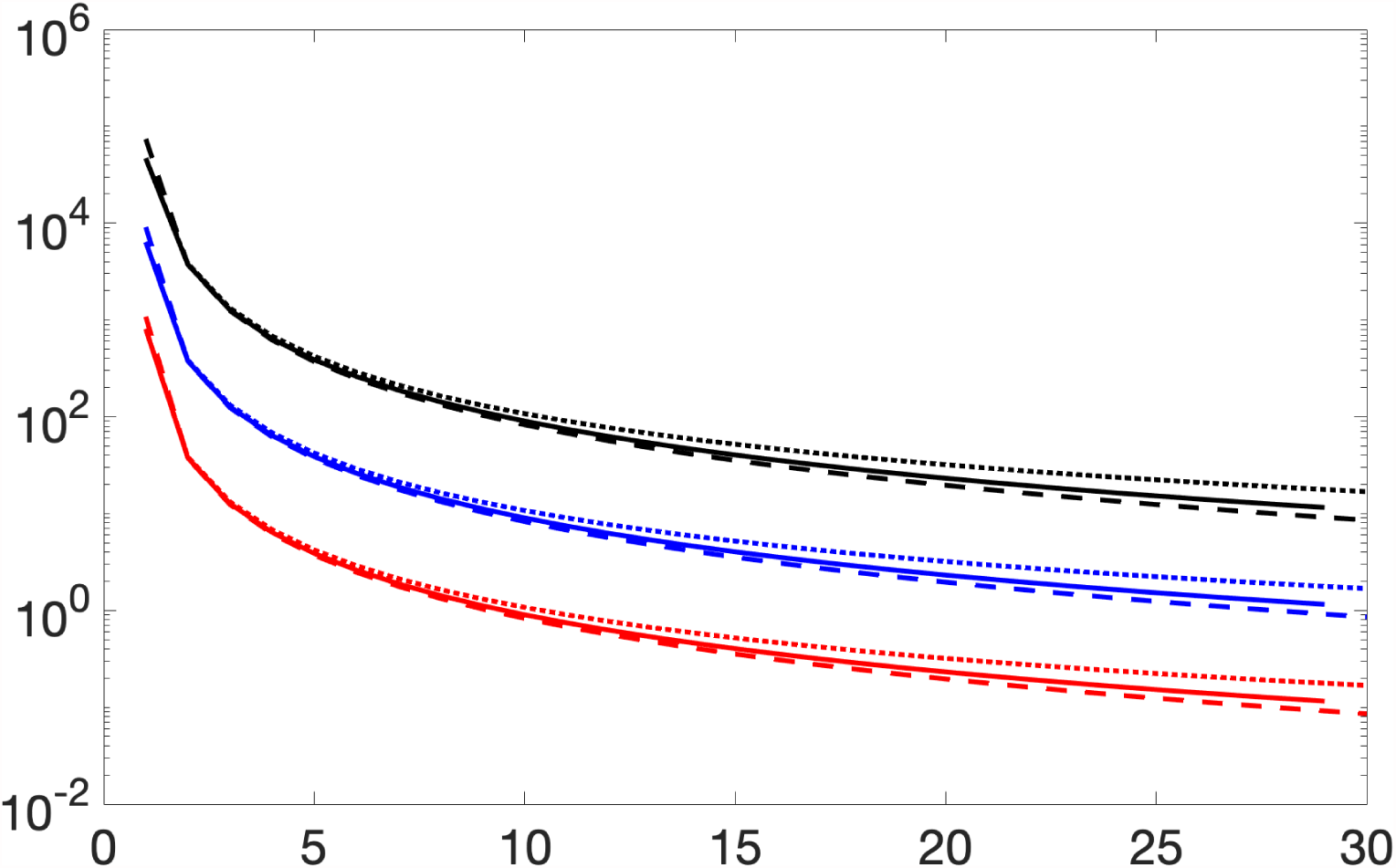
Comparison of expected SFS based on the hypergeometric formula (20) with parameters as in Table 1 (dotted lines), based on Griffiths-Tavaré theory (continuous lines), and Durrett’s approximation (dashed lines). Three cases as in Table 1, fast-growing tumors (red), moderate-growing (blue), and slow growing ones (black) are considered.

**Figure 4:**
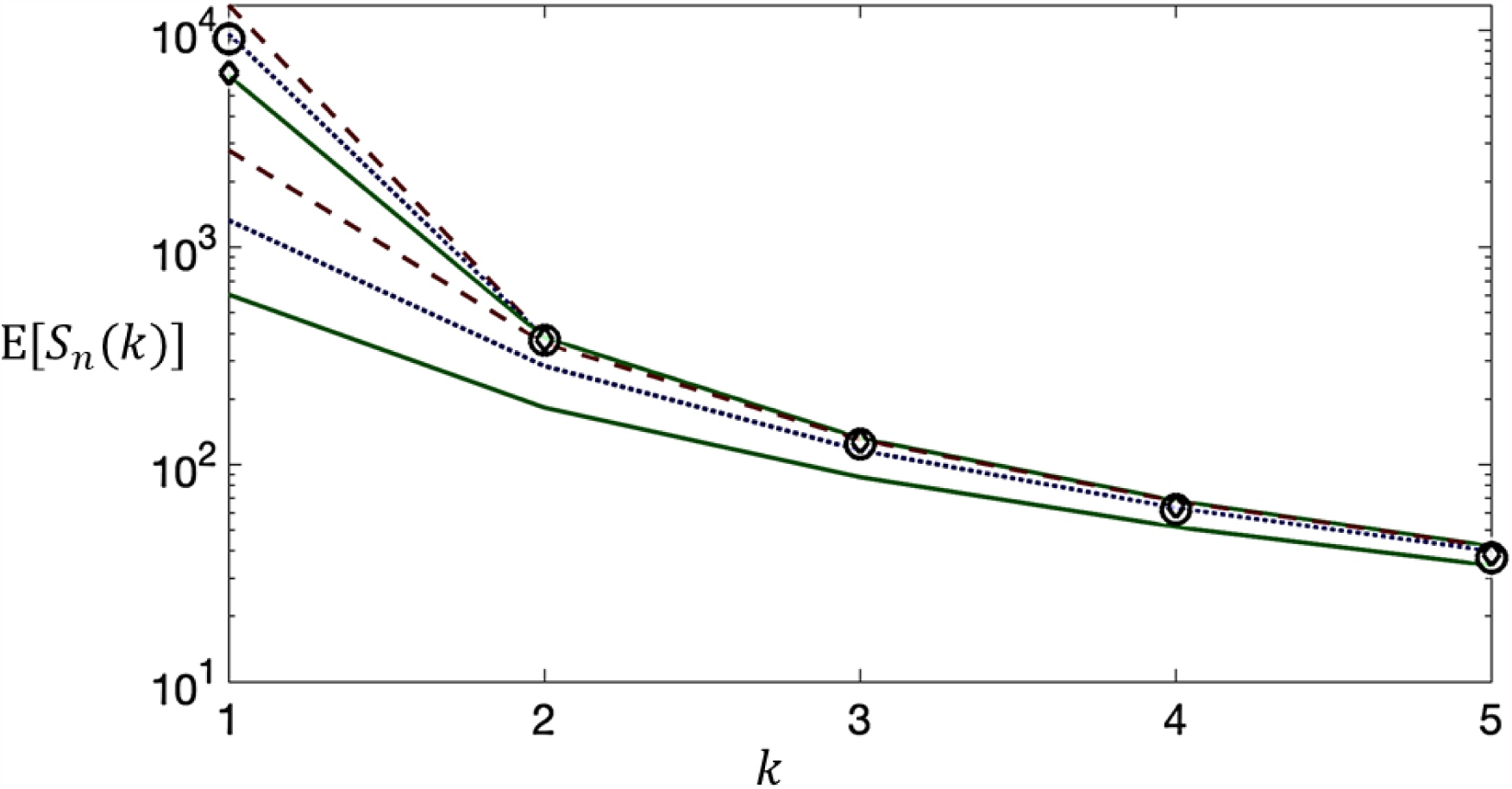
Expected SFS based on the hypergeometric formula (20) with parameters as for the center scenario in Table 1, i.e., *N* = 10^7^, *n* = 30 and *r* = 0.04029, but with 1 *− α* = 10^*−*8^, 10^*−*6^, 0.0001, 0.01, 0.1, 0.5 (dashed, dotted, continuous, and again dashed, dotted and continuous lines), compared to GT SFS (diamonds) and Durrett approximation (circles) with matching parameters.

Several observations can be made. The hypergeometric spectrum for non-singletons preserves signal (however faint) from 1 *− α* = *bp/r* in addition to the signal from *θ/r*. The hypergeometric spectrum has different tails from the Durrett’s approximation of the coalescent spectrum, although whether these can be distinguished in noisy data set seems quite doubtful. Comparison is further complicated by somewhat different sampling philosophies in coalescent and lbdp approaches. An interesting question is how to apply the fitted theoretical spectra to estimate the growth parameters and particularly the time elapsed from the cell initiating tumor growth (more generally, from the population ancestral individual)? The difficulty becomes clear upon inspection of the asymptotic formula (5). None of the terms depends on *N*, the present-time population size, except for the singleton term 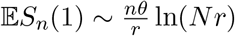, which is equal to *nθt* under exponential growth. However, in genome data singletons are usually indistinguishable from sequencing errors and are therefore discarded. Other terms may be used to estimate the reduced mutation rate *θ/r*.

## 4 Modeling mutation, growth, and selective sweeps

We begin with a simple model for the clonal evolution of a tumor. Imagine that at some time labeled *t*_0_ = 0, the initial malignant cell population (clone 0) arises, grows deterministically in size at rate *r*_0_, these cells acquiring mutations at the rate *θ*_0_ per time unit per genome site. At time *t*_1_ > 0, a secondary clone (clone 1) arises, which differs from the original clone with respect to growth rate (now equal to *r*_1_) and mutation rate (now equal to *θ*_1_). We call this the “selective event”. The new clone arises on the background of a haplotype already harboring *K* mutations. Finally, at *t*_2_ *> t*_1_ > 0, the tumor is diagnosed and a sample of DNA is made available for sequencing. At that point, it is difficult to distinguish cells arising from the two (or more) clones and the resulting sample represents a mixture of DNA from both. The course of events in this tumor history is depicted in Fig. 5.

**Figure 5:**
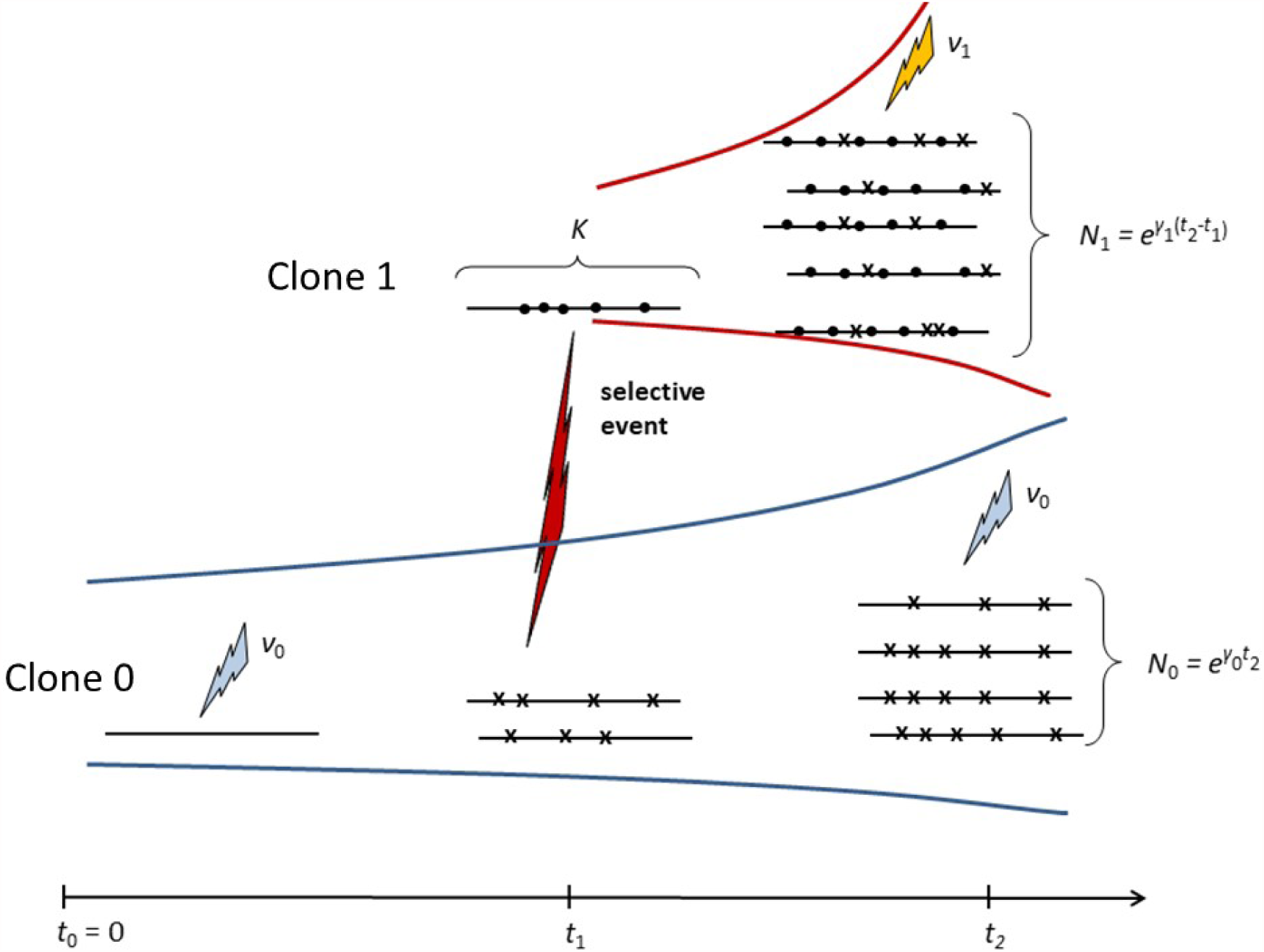
Events in the tumor evolution model. Horizontal intervals denote genomes with mutations denoted as *×*-s. At time *t*_0_ = 0, the initial cell population (clone 0) arises, grows at rate *r*_0_, and mutates at rate *θ*_0_ per time unit per genome site (blue arrows). At time *t*_1_ > 0, a secondary clone 1 arises (red arrow), which grows at rate *r*_1_ and mutates at rate *θ*_1_ (yellow arrows). The new clone arises on the background of a haplotype of *K* mutations (denoted by dots on the genome). At *t*_2_ > *t*_1_ > 0, the tumor is diagnosed and a sample of DNA is sequenced.

**Table 2:**
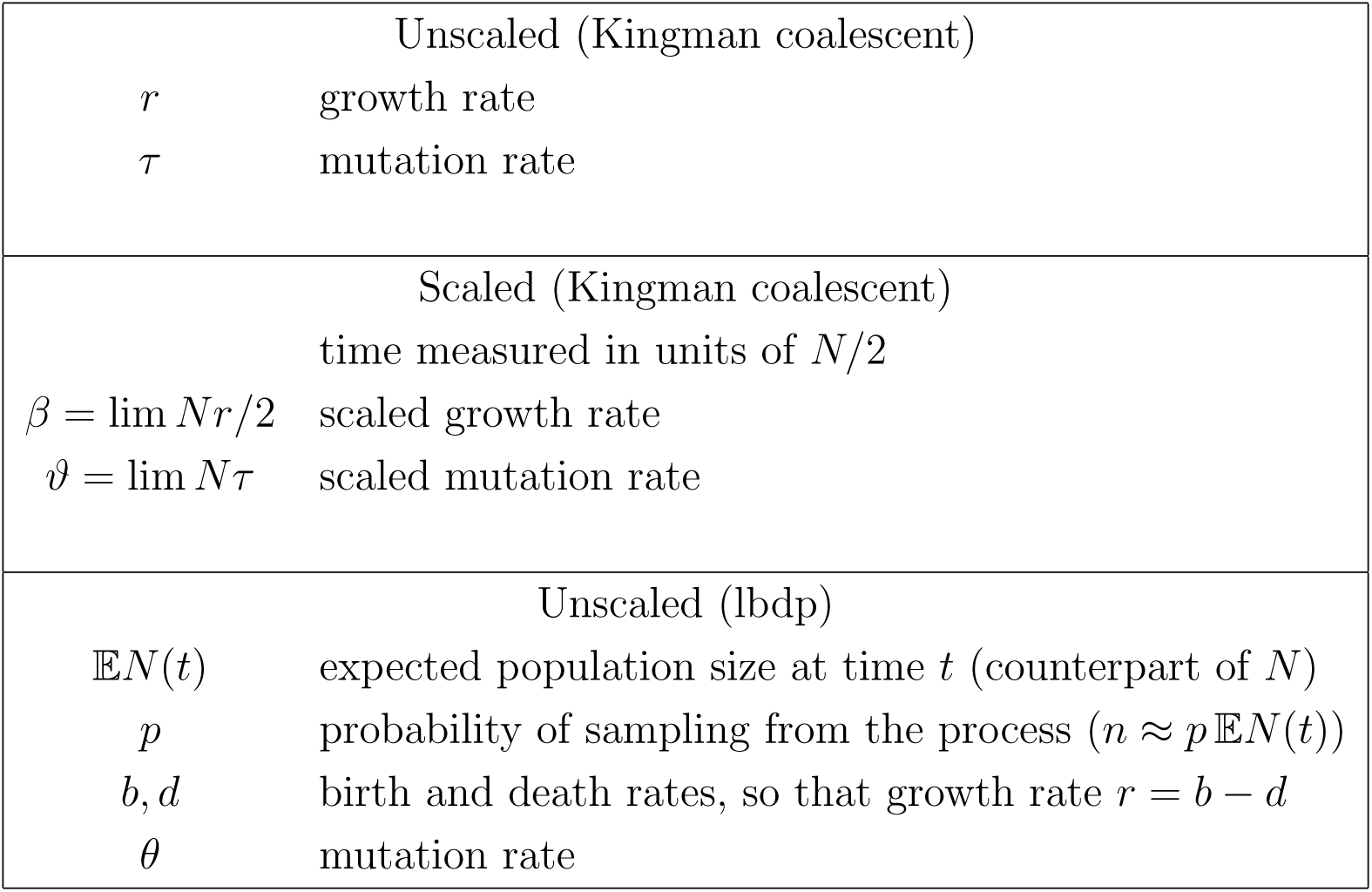
Summary of growth and mutation parameters

We assume that both clones start from single cells, so that the sequenced sample comes from *N* = *N*_0_ + *N*_1_ cells, and the number of cells in each clone is

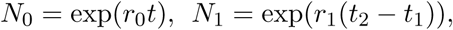

and the fraction of clone *i* cells is approximately equal to

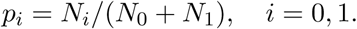

Based on this, we use the SFS from Section 3.1 to estimate the expected site frequency spectra and then compare these to data, to obtain information concerning the natural course of tumor development. As explained before, we use scaled parameters *β*_0_, *β*_1_, *ϑ*_0_, *ϑ*_1_, instead of *r*_0_, *r*_1_, *θ*_0_, *θ*_1_, respectively.

### 4.1 Sampling formulae

We adopt the coalescent model with infinitely-many sites mutation and exponential population growth described in Section 2.1. We take a sample of *n* = *n*_0_ + *n*_1_ cells from the *N* cells in the tumor, *n*_*i*_ coming from clone *i*. We also define

*q*_*n,k*_ = 𝔼*S*_*n*_(*k*), *k* = 2, …, *n*, the expected number of variants present in *k* copies in the sample of *n* sequences

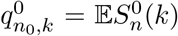, *k* = 2, …, *n*_0_, the expected number of variants present in the *k* copies in a sub-sample of *n*_0_ sequences

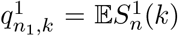, *k* = 2, …, *n*_1_, the expected number of variants present in the *k* copies in a sub-sample of *n*_1_ sequences

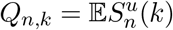, *k* = 2, …, *n*, the expected number of variants present in *k* copies in the sample of *n* sequences from the union of clone populations 0 and 1

We use the approximate version of the expression for the *q*_*n,k*_, given in (5),

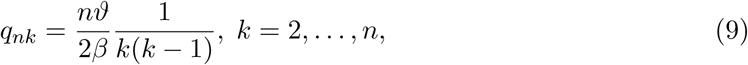

and we ignore singletons.

If we knew *n*_0_ and *n*_1_, the expected number of variant sites represented *k* times in the sample would be 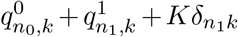, where *δ*_*lk*_ = 1 if *l* = *k*; = 0 if *l* ≠ *k*. However, if each of the *n* cells is randomly chosen from the two sub-clones, then (*n*_0_, *n*_1_) is a random draw from the multinomial distribution, i.e.

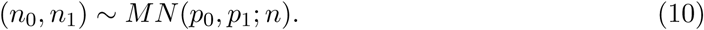

Therefore the expected count of variants present in *k* copies in the sample of *n* cells is

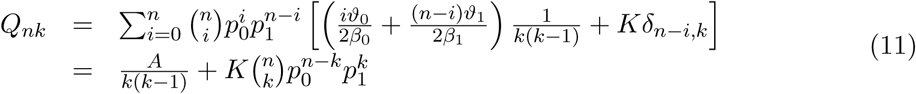

for *k* = 2, …, *n*, where

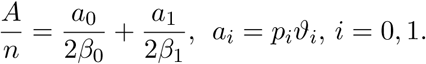

The model can be generalized to the case of *H* more clones arising at different times. The previous expression now assumes the form

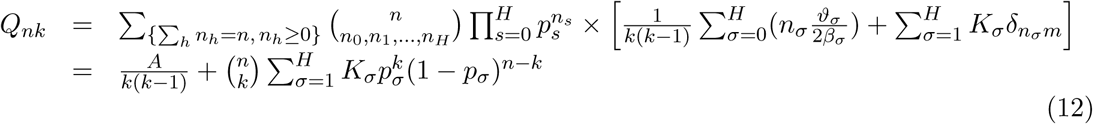

where the notation is analogous to the two-clone case in (11). We will use these expressions, taking into account the sampling effects, in Section 6.

### 4.2 Model parameters and their interpretation

We return to the two-clone case. Equation (11) can be represented in the following form

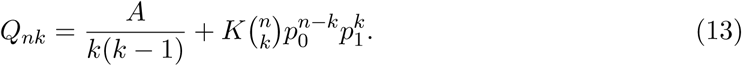

Given SFS data, and the value of *n*, we are able to obtain an optimal least-squares fit by varying three parameters:

*A*, proportional to the mass of the spectra corresponding to the intervals before and after the selective event;

*p*_1_ = 1 *− p*_0_, the fraction of cells in sub-clone 1; and

*K*, the number of variant sites constituting the background haplotype of the selective event.

The parameters listed above are functions of the intrinsic parameters of the model: times *t*_1_ and *t*_2_, growth rates *β*_0_ and *β*_1_ and mutation rates *ϑ*_0_ and *ϑ*_1_. Some of their values can be constrained, based on additional information available in part of the TCGA data.

## 5 Introduction to genome sequencing

The previous sections have introduced our modeling framework, and we now need to make the connection between this and the sequencing data we will exploit. To this end, we provide a brief introduction to technical issues related to genome sequencing. These will be helpful in understanding of sampling issues described in Section 6, and the interpretation of the resulting analysis.

### 5.1 Principles of genome sequencing

Genome sequencing is the methodology used to obtain the nucleotide sequence of DNA in biological cells. Most genome sequencing techniques are based on fragmentation of the DNA extracted from a specimen into shorter fragments (usually 300 - 800 nucleotides in length). So-called Next Generation Sequencing allows massively parallel sequencing reactions which, while focusing only on small sequence fragments, result in hundreds of gigabases of sequencing reads in a single run. This significantly speeds up the process and reduces costs, however it also has some drawbacks. In order to cover the entire sequence of interest many more fragments are needed than determined by the length of the sequence divided by the fragment length. This is caused by the random sampling process of fragments, which provides uneven spread if the number of fragments is low.

Sequencing of short sequence fragments, termed reads, is followed by assembly, which combines them into a collection of single contiguous letters termed contigs. This can be achieved using a human reference sequence, to which the reads are aligned with some leeway allowing for detection of sequence variants not conforming to the reference.

### 5.2 Reads and coverage

Sequencing coverage, also known as sequencing depth, refers to the total number of unique reads covering a particular position of genomic sequence. The higher the coverage, the more precise are the experimental outcomes, but the cost of the experiment also increases significantly. The mean coverage is therefore usually selected based on the goal of the experiment and available funding. By knowing the total size of the library to be sequenced, the error rate of the sequencing instrument and the methodology used, it is possible to predict the coverage and adjust it in a variety of ways.

Different experimental strategies require different coverage levels. For example, in order to detect copy number changes a mean coverage of 1x is sufficient in whole genome sequencing. However, for variant detection the value should be much higher, reaching a minimum recommended value of 15x for germline variants, 30x for somatic cancer associated variants (Bentley et al. (2008)) and 60x for indel detection (Fang et al. (2014)). Those values should be even higher in case of exome sequencing due to very uneven coverage levels (Meynert et al. (2013)). Expected and attained coverage levels might differ significantly since some of the reads can be lost due to effects such as low read quality, inability to align the read to the reference and high read duplication level.

### 5.3 Genome variability: SNV, CNV and other

Genomic variability is identified by either studying changes in the coverage level, as in the case of copy number variation (CNV) or changes in the nucleotide sequence compared to the reference genome. Significant increase in the number of reads at a particular chromosomal region indicates copy number gain while a decrease is associated with copy number loss. This process can also be observed for an entire chromosome in a case of chromosomal monosomy or polysomy, for example three chromosomes might be observed instead of the normal two, such as in the case of Down syndrome, which is caused by a trisomy of chromosome 21.

Changes in the nucleotide sequence are usually harder to identify since they have to be distinguished from sequencing errors caused, for example, by incorrect base calls or polymerase slippage (Viguera et al. (2001)). High coverage is therefore important for variant detection since the confidence of a variant call increases with the number of observed reads that show them. The variant allele frequency (VAF) is the number of reads with a particular variant, which can be either a single nucleotide variant (SNV) or an indel (insertion/deletion). Additionally, by comparing tumor and normal cell samples it is possible to differentiate germline from somatic changes, the latter being often the main contributing factor to development of the tumor. Zygosity (ploidy) can be determined by studying variant allele frequency. In the case of germline variants this usually reflects the number of variant alleles. A variant allele frequency of 0.5 indicates heterozygosity while a frequency of 1 indicates homozygosity. This is much more complex for cancer cells, which are often a mixture of cells with various genotypes. A variant allele frequency of 0.5 might indicate heterozygosity, however it is also possible that 50% of the sequenced cells are homozygous for that variant while the remaining fraction doesn’t show any variant. This becomes much more complicated for values other than 0.5 and 1 and for regions with abnormal copy numbers. A variant allele frequency of 1 might not indicate that the variant is present in both chromosome copies, but can also be observed if the variant exists in one allele while the second allele is missing due to copy number loss. This event is known as loss of heterozygosity (LOH).

### 5.4 Tumor genomes

Cancer studies that aim at identifying genomic variants require high coverage levels due to the fact that a tumor is usually composed of cells with significantly different genotypes. Detection of rare variants, found only in a small fraction of cells, requires even higher coverage. This is typical of experiments which aim at studying cancer evolution.

Increased coverage significantly improves the reproducibility of the experiment, allowing more precise identification of true variants from sequencing errors. However in cancer studies the problem is much more complicated. Due to sampling, rare variants specific to a small fraction of cells might be detected only in one of the replicates, however the overlap might not increase with an increased coverage level since this will lead to the discovery of new, even rarer, variants that might be detected only in one of the replicated samples (Jaksik et al. (2018)).

Mutations may appear anywhere in the genome and despite the fact that some regions and positions can mutate more often, most mutations are believed to have no impact on the phenotype. Only a subset of mutations lead to cancer progression and it is believed that there are several genes that need to be altered by mutations, indels or CNV in order for cancer to progress. Those that need to be inactivated are known as tumor suppressor genes, which are believed to protect the cells from carcinogenesis. Another class includes oncogenes that need to be over-activated by mutations. Mutations that drive cancer progression are known as *drivers* while those that do not have a direct impact are called *passengers*. For a more clinical view of whole-genome sequencing for identification of targetable variants in cancer, see Wrzeszczynski et al. (2018) for example.

## 6 Sampling from the SFS

One of the conceptual problems with using the model-based expectations of the site frequency spectrum is how to take into account the sampling process. Indeed, the empirical SFS are not based on the cell population, but on DNA reads (fragments) sampled from the genomes of the cells. Therefore, it is necessary to proceed with care. Under simplifying assumptions, we can obtain unbiased estimates of the expected SFS, given a parametric model of either coalescent or lbdp type. The assumptions are as follows:

1. DNA fragments (reads) used to estimate variant allele frequencies (VAF) originate from a population of cells, with variant genomes representative of a given tumor or a portion of the tumor.
2. For each particular mutation site, each read covering this site originates from a different cell. This seems to be a reasonable assumption, as the number of such reads is usually at most of order 10^2^, while there are around at least 3-5 orders of magnitude more tumor cells in a cubic millimeter of tumor tissue (Del Monte 2009).
3. For a given mutation site, the numbers of reads covering it is considered a random variable (generically named *R*) drawn from a distribution which does not depend on the site position in the genome. This assumption can be relaxed in a variety of ways, but it is used here for simplicity. The distribution of *R* is estimated from coverage data.
4. For a given mutation site, given coverage *R*, the count *Z* of variant reads has a binomial distribution Binomial (*R, ϕ*), where *ϕ* is the relative frequency of this mutation among the tumor cells.

Unfortunately, it seems difficult to exploit the higher moments of the SFS, as this requires using mixed moments of variant counts at different sites. The papers by Sargsyan (2015) and Klassman and Ferretti (2017) lay out the necessary theory, which is however quite complex.

### 6.1 Binomial sampling and data pre-processing

In the following subsections we develop estimates of the coalescent SFS based on binomial sampling. Since various types of thresholds might be used to pre-process genome data, we would like the transformations to robustly reproduce the effects of varying the thresholds, while keeping constant the parameters, such as *A*, *K*, and *p*_1_, of the underlying model. We will see that in some instances this works on real-life tumor data with some precision, while in some others it does not.

#### 6.1.1 Sampling

Let *n* be the total number of cells at the bottom of the tree (that is, all the cells in the tumor sample). The model-based expected SFS is the sequence 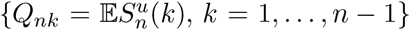, i.e. the expected number of mutations that occur in exactly *k* out of these *n* cells (see (11)). For the *i*th mutation of the 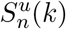 occurring in *k* cells, the probability mass function (pmf) of read coverage is 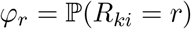, *r* = 1,2, …, and the number of cells with mutation *i* in the sample is *Z*_*ki*_, where conditionally on *R*_*ki*_, we assume a binomial distribution with probability of success *k/n* and *R*_*ki*_ trials (see hypothesis 4 earlier on):

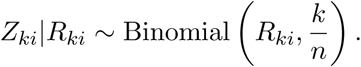

Relative frequency of this particular mutation in the sample is *Z*_*ki*_/*R*_*ki*_.

For 0 ≤ *x*_1_ < *x*_2_ ≤ 1, we are interested in the expectation of Ω(*x*_1_, *x*_2_), the number of mutations with sampling frequencies within (*x*_1_, *x*_2_]:

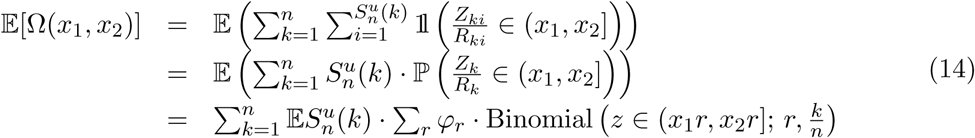

where Binomial 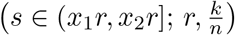 is the probability that random variable *s* with distribution Binomial 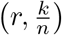 belongs to the interval (*x*_1_*r, x*_2_*r*]. Given a theoretical SFS (for instance given by (11)), we can then compute the expected SFS, taking into account sampling effects, by partitioning 0 = *x*_1_ < *x*_2_ < … < *x*_*K*_ = 1 and applying (14) for each interval.

In general, the read coverage pmf {*ϕ*_*r*_} varies among patients and tumors. In our computations, we use a “personalized” estimate of the coverage distribution, which is based on the tally of reads for all sites in each sample and is usually available from sequencing data.

#### 6.1.2 Pre-processing and its influence on SFS visualization

Mutations with small frequencies may be difficult to distinguish from technical errors. Data are therefore usually pre-processed before further analysis. Specifically, it is a usual practice to remove from genome statistics variants that are present in only few reads, since these may be confused with sequencing errors. A procedure of this kind has been proposed among others by Williams et al. (2018), who disregard variants present in less than five reads. We slightly generalize this approach.

We consider two pre-processing schemes:

1. Disregard mutations with fewer than *L* variant reads. This means that the new variant read count 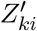 is such that 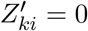 with probability 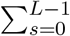 Binomial (*s*; *r*, *k/n*) or 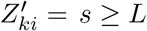 *s* ≥ *L* with respective probabilities Binomial (*s*; *r, k/n*).
2. Disregard mutations with fewer than *M* total read coverage. This alters the coverage pmf {*ϕ*_*r*_}.

We note that different pairs (*L, M*), mask differently the neutral and selective (“hump”) components of the SFS. The following are some interesting cases:

- *L* > 1 makes singletons invisible.
- In general, larger values of *L* make it difficult to visualize the existence of the neutral clone represented by the descending component at the left end of the SFS, and therefore low *Z/R* ratios.
- However, large values of *M* with moderate *L* may allow uncovering of the neutral component, since then more variants with low *Z/R* ratios may be visible. The limitation is that there are enough variants with high *R* values.

In the next section, we will show on biological examples how this may work.

### 6.2 Examples

We now study the effects of pre-processing on the SFS from patients from The Cancer Genome Atlas (TCGA) collection. The parameters from fitting the SFS are shown in Table 3. The numbers show an interesting trend, which may have relevance for estimation of total mutation count in the tumor sample (see Discussion).

**Table 3:** Parameters from fitting the SFS in the TCGA collection. For every case, the number of mutations reported from the sequencing data is shown, as well as the number of humps in the fitted SFS, and parameters *A* and (*K, p*) for each hump (Equation (13)).

| Cases | Total number of mutations | Number of humps | A | K | p |
| --- | --- | --- | --- | --- | --- |
| TCGA-AA-3977 | 1,051,861 | 2 | 20,000,000 | 200,000 | 0.23 |
|  |  |  |  | 450,000 | 0.35 |
| TCGA-A6-6141 | 254,759 | 2 | 2,600,000 | 42,000 | 0.24 |
|  |  |  |  | 180,000 | 0.56 |
| TCGA-86-A4D0 | 1,104 | 4 | 15,000 | 140 | 0.31 |
|  |  |  |  | 280 | 0.44 |
|  |  |  |  | 80 | 0.61 |
|  |  |  |  | 50 | 0.78 |
| TCGA-62-A46O | 2,854 | 5 | 15,000 | 400 | 0.27 |
|  |  |  |  | 800 | 0.41 |
|  |  |  |  | 700 | 0.58 |
|  |  |  |  | 200 | 0.80 |
|  |  |  |  | 150 | 0.91 |

#### 6.2.1 Colon cancers with polymerase *ε* mutator phenotype

We start with two cases displaying the Polymerase *ε* mutator phenotype, which results in a very large number of mutations caused by proofreading errors of DNA replication due to faulty polymerase. These are most frequently colon cancers. Naturally, in these cases, sites have unusually high coverage and therefore using a high threshold *M* does not remove all information from the sample.

Both cases have been pre-processed using four different thresholds of *L* = 5, 10, 15, and 20. The theoretical SFS based on expressions (14) and (12) is fitted to the patient’s data with threshold *L* = 10. The resulting parameter set (consisting of *A* for the neutral slope, and (*K*_*σ*_, *p*_*σ*_) for each hump, with *σ* = 1, *…, H*) is then used to compute the sampled SFS for the other thresholds (*L* = 5, 15, 20) and compare them with the correspondingly thresholded data. We also examine the effects of thresholding the total read counts. We consider four different thresholds, *M* = 20, 30, 40, and 50, all with *L* = 5. Results are shown in Figures 6 and 7.

**Figure 6:**
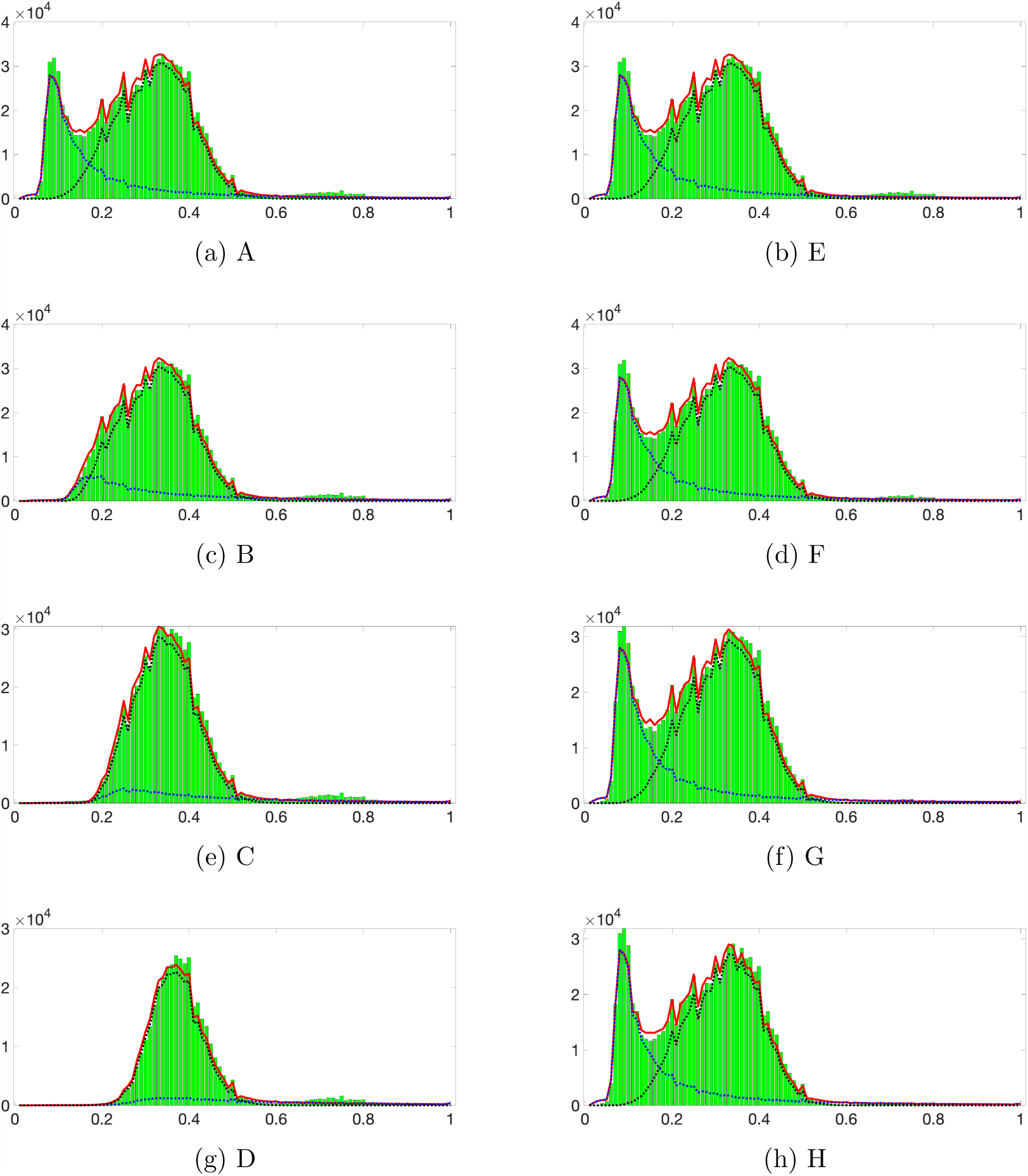
Fitting the SFS of case TCGA-AA-3977 (colon cancer). The theoretical SFS (red lines, Equation (14)) is fitted to the patient’s SFS (green bars). The blue and black dotted lines denote the contribution of the neutral part and binomial humps in the fitted SFS, respectively. Threshold combinations of variant and total read counts: [A]: *L* = 5, *M* = 0, [B]: *L* = 10, *M* = 0, [C]: *L* = 15, *M* = 0, [D]: *L* = 20, *M* = 0, [E]: *L* = 5, *M* = 20, [F]: *L* = 5, *M* = 30, [G]: *L* = 5, *M* = 40, [H]: *L* = 5, *M* = 50.

**Figure 7:**
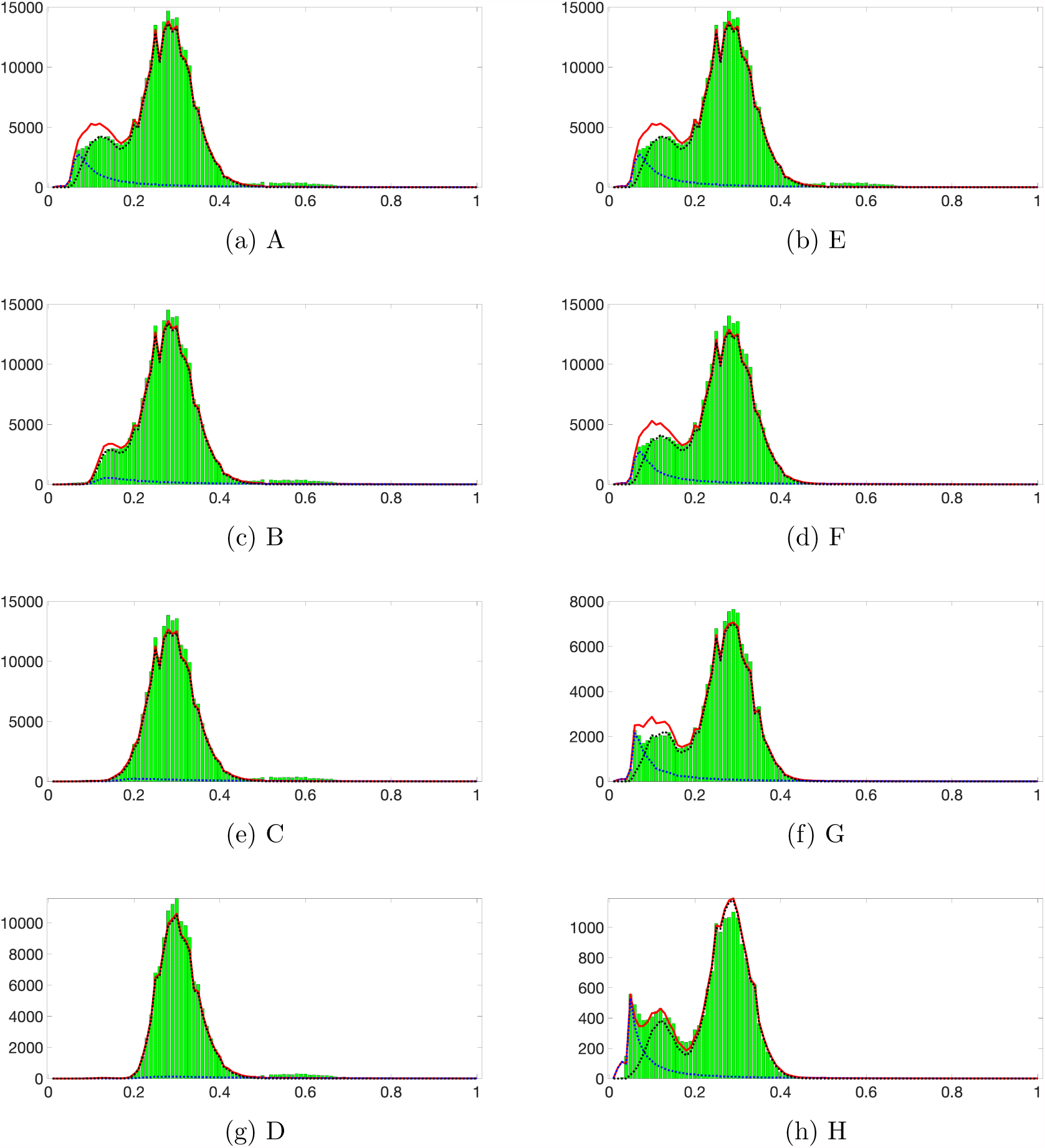
Fitting the SFS of case TCGA-A6-6141 (colon cancer). The theoretical SFS (red lines, Equation (14)) is fitted to the patient’s SFS (green bars). The blue and black dotted lines denote the contribution of the neutral part and binomial humps in the fitted SFS, respectively. Threshold combinations of variant and total read counts: [A]: *L* = 5, *M* = 0, [B]: *L* = 10, *M* = 0, [C]: *L* = 15, *M* = 0, [D]: *L* = 20, *M* = 0, [E]: *L* = 5, *M* = 20, [F]: *L* = 5, *M* = 50, [G]: *L* = 5, *M* = 80, [H]: *L* = 5, *M* = 100.

For the case of TCGA-AA-3977 (colon cancer), the SFS for *L* = 10 can be well fitted with the theoretical expression (Figure 6-B). Moreover, the resulting parameter set accurately recreates the SFS with the other thresholds. Even the pattern of the fluctuations within each SFS can be reproduced, likely because data-based coverage distribution has been used. This reinforces the relevance of the theoretical model and the sampling scheme, including the pre-processing step. It can also be observed that varying the conditioning thresholds for variant and total read counts leads to very different visualizations.

Although higher thresholds *L* result in more reliable SFS (as false positives due to technical errors are less likely), they also gradually dissolve the neutral part of the spectrum that dominates the region with low VAF. This neutral slope can be easily recognized at *L* = 5 (Figure 6-A, 8% mutations in the dataset are discarded) but at *L* = 20, only the hump can be observed (Figure 6-D, 63% mutations are discarded).

On the other hand, increasing the threshold *M* preserves the overall structure of the SFS. Comparing the SFS with *L* = 5, *M* = 20 (Figure 6-E, 8% mutations in the dataset discarded) and that with *L* = 5, *M* = 50 (Figure 6-H, 21% mutations discarded), we observe a slight decrease in the height of the hump, while the neutral slope remains intact.

The results for case TCGA-A6-6141 (colon cancer), however, show a somewhat different picture. The SFS for various thresholds of *L* and *M* reveals a small hump at low frequencies (*f <* 0.2), which can be interpreted as the neutral slope. Fitting the SFS under this assumption for *L* = 10 results in a good fit, consisting of the neutral slope and one hump at VAF *f* = 0.28, and can recreate the SFS for higher *L*, but presents a discrepancy for *L* = 5, with or without additional conditioning on *M*. At very high total read coverage (*L* = 5, *M* = 100), we can hypothesize the reason for this inconsistency: the low frequencies area seems to contain one small hump close to the neutral slope, and this combination gets blurred at more relaxed conditioning. This small hump may result from neutral sweep(s) during the tumor evolution. Under the assumption of a second hump, the fits conform to the clinical data for all conditioning on *L* and *M* (Figure 7).

#### 6.2.2 Lung cancer

Two TCGA samples from lung cancer were also fitted to the model (Figures 8 and 9). There are several differences between these samples and the cases displaying the Polymerase *ε* mutator phenotype. First, the data result from whole-exome sequencing, which only reports mutations in the protein-coding regions of genes, while the previous cases resulted from whole-genome sequencing, which reported mutations in the non-coding regions as well. This contributes, along with absence of Polymerase *ε* mutation, to the number of mutations in these cases being much lower than in the mutator dataset, and SFS being accordingly more noisy.

**Figure 8:**
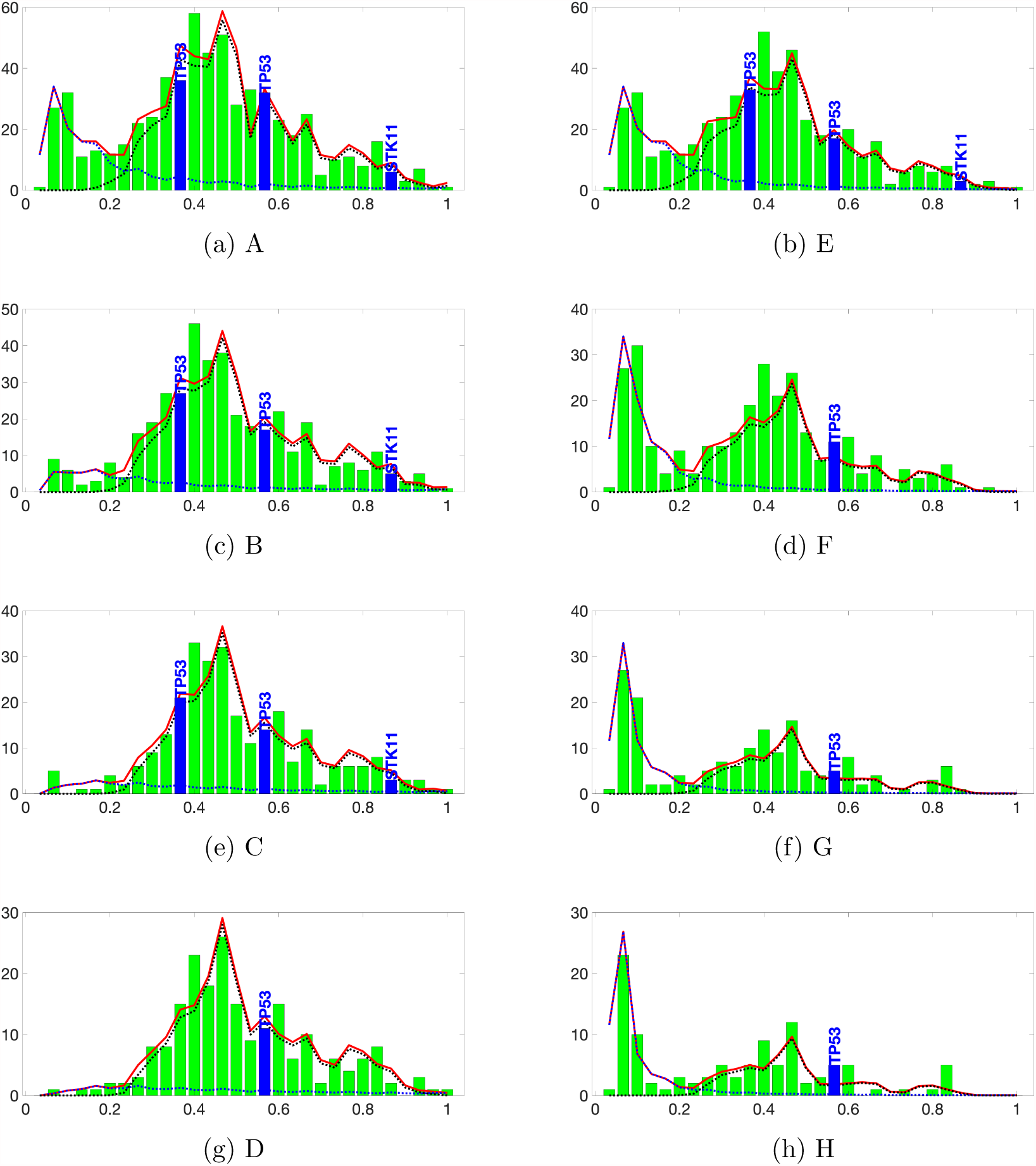
Fitting the SFS of case TCGA-86-A4D0 (lung cancer). The theoretical SFS (red lines, Equation (14)) is fitted to the patient’s SFS (green bars). The blue and black dotted lines denote the contribution of the neutral part and binomial humps in the fitted SFS, respectively. Driver mutations are denoted in blue at their frequencies. Threshold combinations of variant and total read counts: [A]: *L* = 5, *M* = 0. [B]: *L* = 10, *M* = 0. [C]: *L* = 15, *M* = 0. [D]: *L* = 20, *M* = 0. [E]: *L* = 5, *M* = 20. [F]: *L* = 5, *M* = 50. [G]: *L* = 5, *M* = 80. [H]: *L* = 5, *M* = 100.

**Figure 9:**
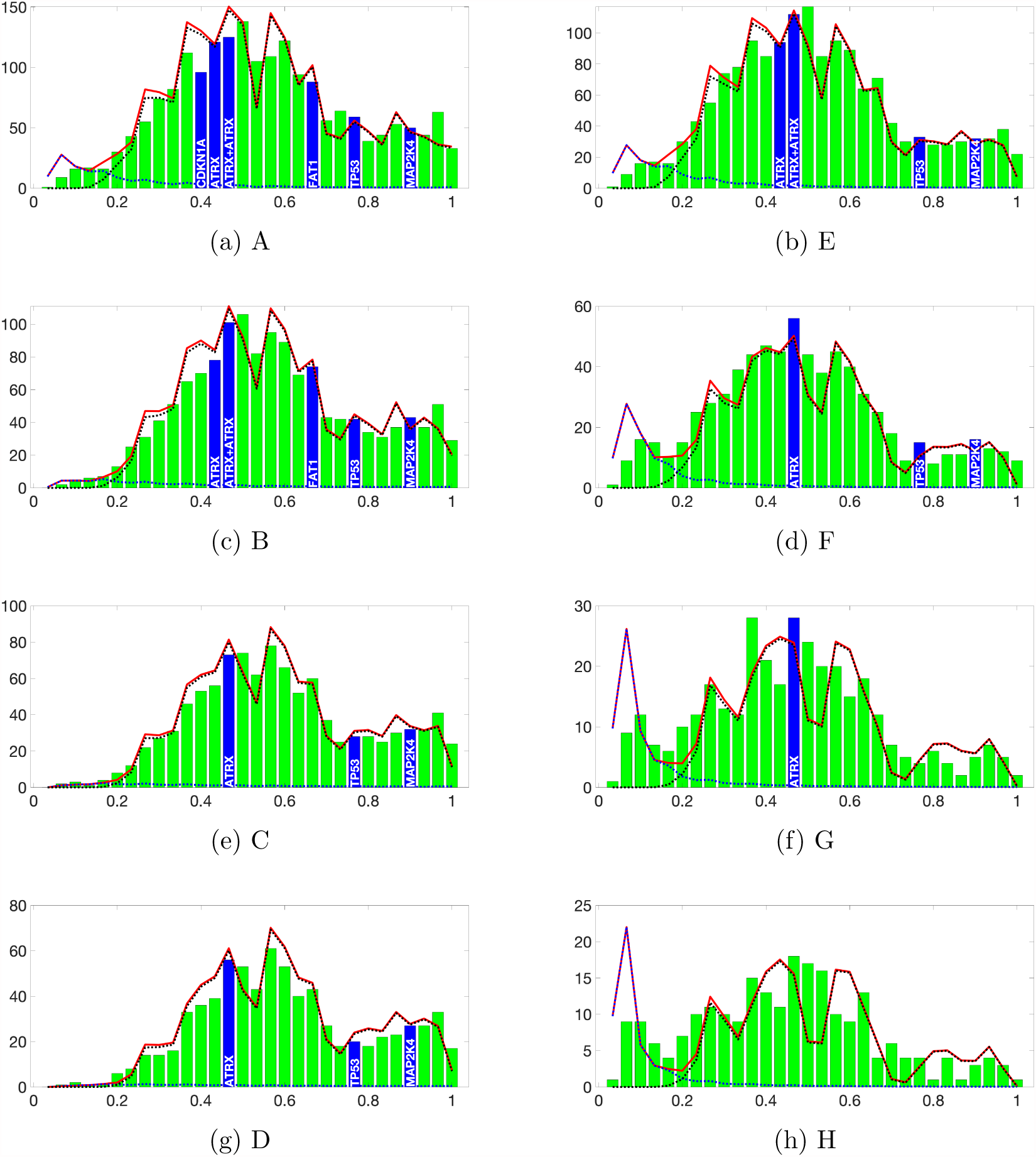
Fitting the SFS of case TCGA-62-A46O (lung cancer). The theoretical SFS (red lines, Equation (14)) is fitted to the patient’s SFS (green bars). The blue and black dotted lines denote the contribution of the neutral part and binomial humps in the fitted SFS, respectively. Driver mutations are denoted in blue at their frequencies. Threshold combinations of variant and total read counts: [A]: *L* = 5, *M* = 0, [B]: *L* = 10, *M* = 0, [C]: *L* = 15, *M* = 0, [D]: *L* = 20, *M* = 0, [E]: *L* = 5, *M* = 20, [F]: *L* = 5, *M* = 50, [G]: *L* = 5, *M* = 80, [H]: *L* = 5, *M* = 100.

Figure 8 shows the results of fitting the case TCGA-86-A4D0. While the two TCGA-WGS-KEEP cases can be fitted with two humps, this case is fitted with four humps. This is in agreement with the existence of various driver mutations at different frequencies. Each of the drivers, therefore, could be associated with one or more humps in the fitted SFS. The difference between the real and fitted SFS is most severe at frequency *f ≤* 1*/*30. This may be because the mutations at low frequencies are more likely to be disregarded, as they can be confused with technical errors. However, the fitted SFS is in overall agreement with the data across different thresholds of *L* and *M*.

The results of fitting the TCGA-62-A46O case (Figure 9) share two signature aspects with the case TCGA-86-A4D0 discussed above. First, the SFS can only be fitted with five humps. Again, the data shows that there are multiple driver mutations in this sample at various frequencies, which may support the high number of humps. Second, the fitted SFS has a peak at low frequencies, which is not supported by the data. On the other hand, this case shows an interesting phenomenon: the fitted SFS is good for various thresholds for *L* (Figure 9-A, B, C, D) but becomes worse as *M* increases (Figure 9-E, F, G, H). One aspect to consider, however, is that increasing *M* rapidly decreases the number of mutations in the SFS. At *M* = 100, the SFS contains only 234 mutations, out of 2854 mutations in the data (Figure 9-H).

This example and the case TCGA-A6-6141 (Figure 7) showcase the two sides of fitting the SFS at high coverage: it can reveal information that is otherwise hidden at lower coverage, but the low mutation count can make the SFS difficult to fit without a large number of humps.

### 6.3 Linear birth-death process with sweeps

#### 6.3.1 Simulating selective sweeps

It seems useful to compare the SFS with selective sweeps based on the coalescent approach to those based on the lbdp approach. While mathematical results have not been developed in the latter setting, we experimented with simulation code that allows generating in time of the order of minutes a random lbdp tree consisting of 10^4^ or even 10^5^ cells. If the cell count is up to 10^3^, we are able to draw the tree using the same convention of enumeration that was used in Section 3.3.

Cells proliferate according to lbdp with rates *b* and *d*. During its lifetime, each cell gathers neutral mutations according to a Poisson process with intensity *θ*. These mutations are shared by progeny of the cell. At a predetermined time point *s*, the cell with the highest number of neutral mutations among all cells alive, acquires an advantageous mutation. This cell initiates a new lbdp (advantageous clone) with rates *b*^®^ and *d*^®^ chosen so that the growth rate is higher than that in the original process. At the end time *T*, the mutation counts of all live cells from the original process and the advantageous clone are determined. The SFS is determined from the frequencies of all neutral mutations, or from a random sample obtained via binomial sampling.

The neutral mutations are partitioned into three subgroups:

- Background mutations: acquired by the selective founder cell or any of its ancestors. These cells are therefore shared among all selective cells and possibly some neutral cells.
- Foreground mutations: acquired by any selective cell. These can be shared among some selective cells but not by any neutral cells.
- Other mutations: neither of the above.

Figure 10 represents an example of the resulting simulated tree. The neutral cells are shown as blue circles and the advantageous cells as green circles. The neutral cells that are ancestors to the selective founder cell are shown in red. In the plot, dead and live cells are indexed so that all descendants of any given cell are grouped together. Cells alive at the final time *T* are shown as solid symbols.

**Figure 10:**
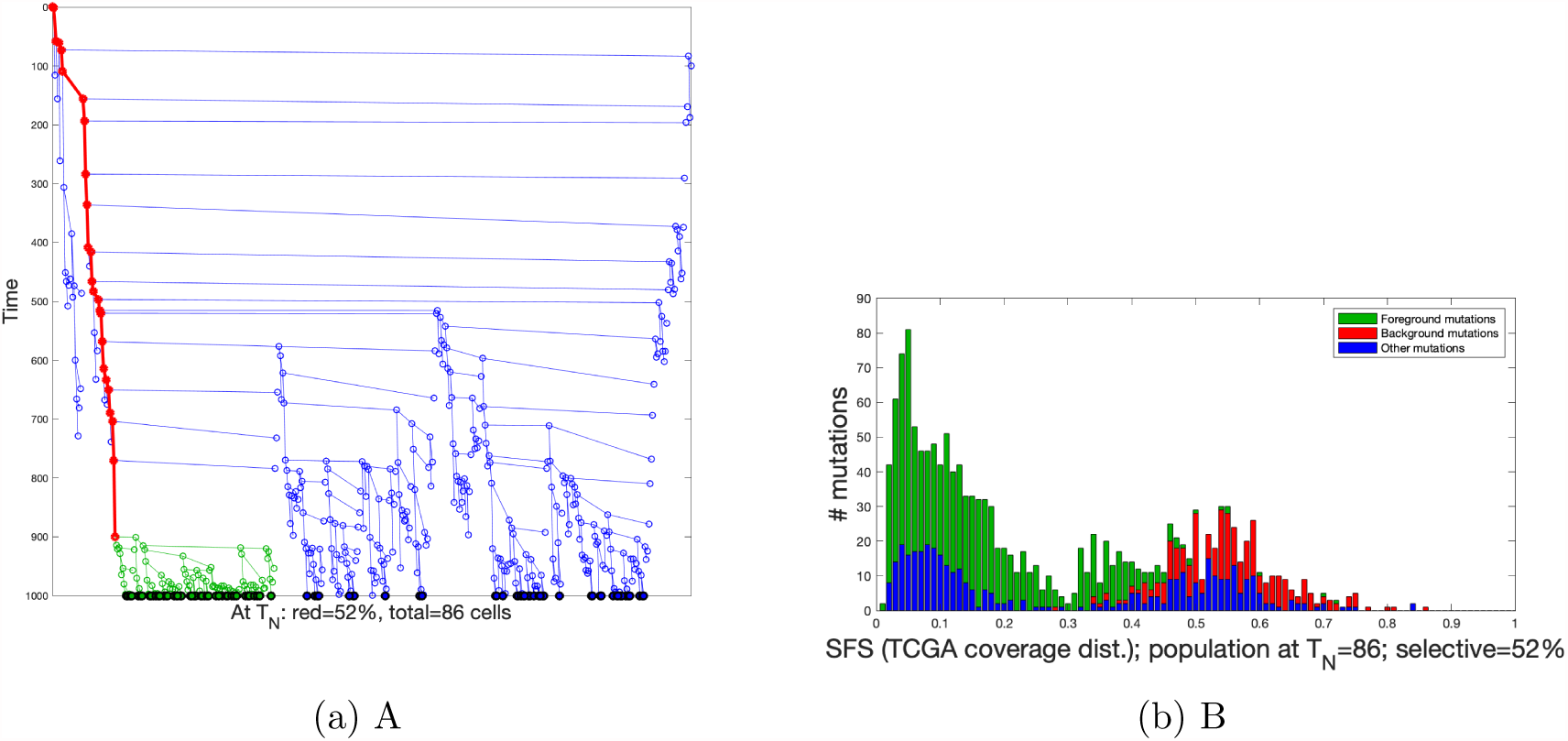
[A]: Example of a simulated tree and resulting SFS. The y-axis is time, x-axis includes invisible indices of cells such that progeny of any given cell is grouped together. The three types of mutations correspond to the SFS. [B]: the SFS resulting from sampling the simulation under the TCGA distribution.

Figure 11, panel A, shows a “trimmed” view of the simulated tree. The cells that have no progeny at final time *T* are removed, since their mutations do not contribute to the SFS. All other aspects are similar to Figure 10.

**Figure 11:**
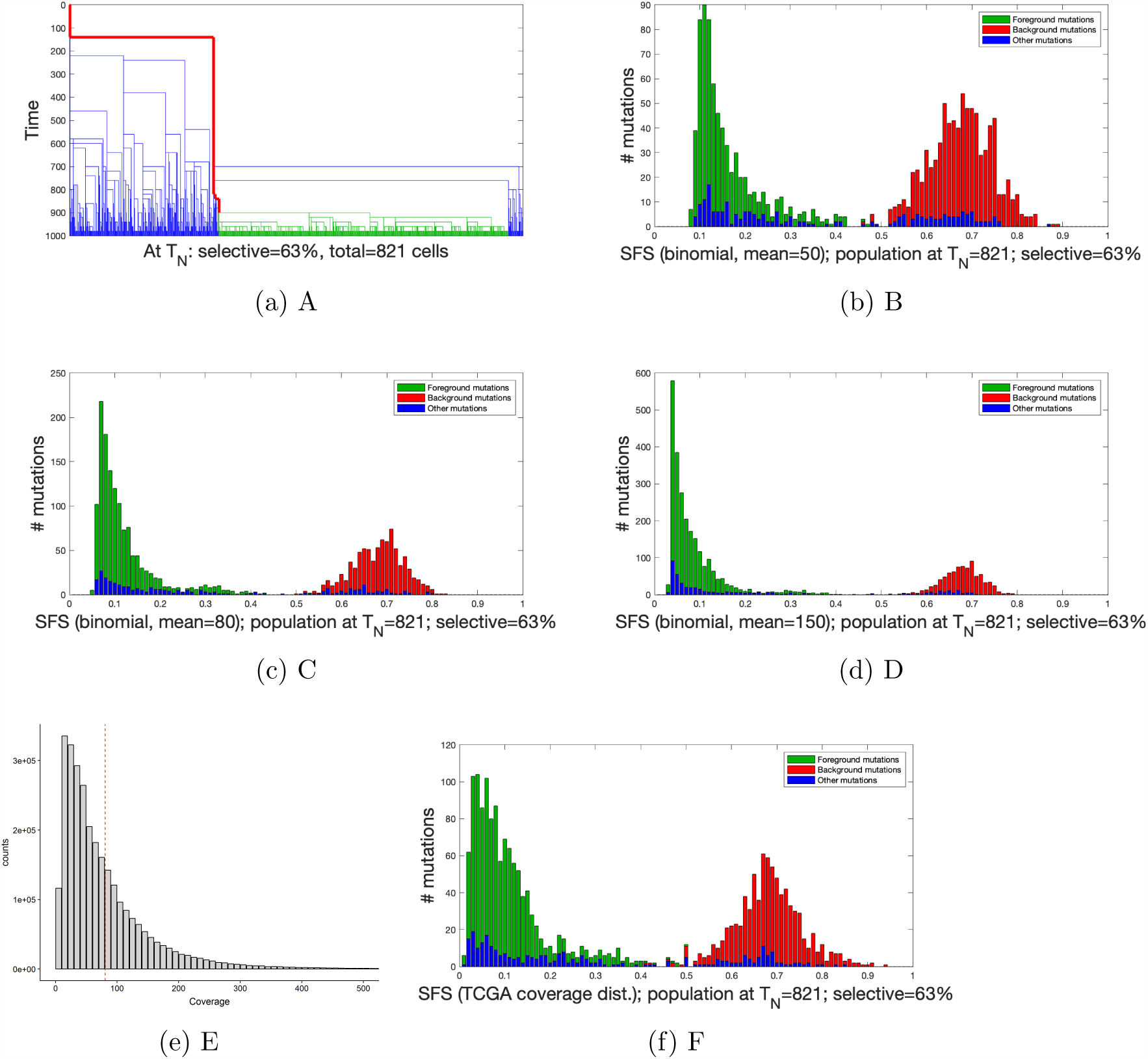
The choice of sampling distribution distorts the resulting SFS. [A]: the simplified presentation of the simulated tree. [B, C, D]: the SFS resulting from sampling the simulation under the binomial distribution with mean 50 (B), 80 (C) and 150 (D). [E]: PDF of the TCGA sampling distribution. [F]: the SFS resulting from sampling from the simulated tree according to the TCGA distribution. Parameters: *T* = 1000, *s* = 800, *b* = 0.0162, *b*^®^ = 0.0721, *d* = *d*^®^ = 0.01, *θ* = 1.

We now discuss how the sampling coverage distribution may affect the SFS from the simulation. A common assumption (e.g. Williams et al. 2018) is that the sizes of samples for detecting mutations follow the binomial distribution. To implement this, we performed one simulation, which resulted in *∼* 10^3^ cells at the final time (Figure 11-A). Four sampling coverage distributions are used: binomial distributions with mean 50, 80 and 150, and the TCGA distribution as in Figure 11, panels B, C, D, and F.

We can observe in Figure 11 that the background mutations form a hump centered around frequency *f* = 0.68, consistent with the fact that the selective clone makes up for 63% of the population. Meanwhile, the foreground mutations form a decreasing slope at low frequencies (*f <* 0.2) which can be explained by the theory in (13). Under deep binomial sampling distributions (Figure 11-C, D), the other mutations also show the characteristic neutral slope, which is more obscure under the TCGA distribution (Figure 11-F).

## 7 Discussion

Our paper has outlined a model-based approach to inferring aspects of the clonal evolution of cancers, using data from the site frequency spectrum of somatic single nucleotide variants found from bulk whole-genome or exome sequencing. We focused primarily on two aspects: stochastic models of tumor evolution adapted from the fields of population genetics and population dynamics, and the effects of “data cleaning” that is often used in the analysis of sequencing data.

The modeling aspects have made a number of simplifying assumptions that make the statistical inference aspects tractable. In the comments below, we address a number of these in more detail.

### Simplicity

Following a review of mathematical models of site frequency spectra based on Kingman’s coalescent and the linear birth-death process, we develop a theory for models of clonal sweeps. We explored its action on simulated and TCGA data-based spectra. The leading principle in our analysis was simplicity. We considered a neutrally evolving cell population, which spawns an advantageous mutant giving rise to a clone with different growth and mutation rates. The clone leads to a hump in the spectrum. More than one such event can be accommodated. This approach allows us to estimate aggregate parameters *A*, *K*, and *p*_1_ of the model. We note that our model is not spatial, nor does it deal with multiregion data explicitly, although extensions are conceivable.

### Simulation approach of Williams et al

Williams et al. (2018) present a simulation-based approach that is based on an lbdp. Our sampling transformation in (14) can be considered an “expected value” version of their data transformation. However, there are notable differences in approach. Williams et al. identify as a separate category the “truncal” mutations, i.e. mutations that arose in the ancestor of the tumor clone. These mutations are present in all tumor cells, however, since they are usually heterozygotic (the other allele being a non-mutant variant), they are present in 50% of DNA strands. Since reads covering a truncal variant site are sampled binomially, truncal mutations form a binomial hump centered at variant allele frequency *x* = 0.5. In contrast, the emerging new selective clone leaves another binomial hump, being a signature of the mutations accumulated in its ancestral cell (which might be called truncal mutations of this particular clone). This hump is centered at VAF *x* ≠ 0.5, depending on the fraction of tumor cells in the new clone. In our experience, the “solitary” humps seem to be rarely centered at VAF *x* = 0.5. This might be a result of contamination. However, please see the discussion of evolutionary history further on. We can easily accommodate truncal mutations by adding an extra hump in Equ. (12).

### Ploidy

In Section 6.1 we treated all tumors as haploid (i.e. with ploidy equal to 1, or with a single copy of each chromosome per cell). This clearly not accurate, as human tumors are usually derived from diploid (ploidy equal to 2) cells. Can tumor cell ploidy be accomodated in our framework? If tumor cells are diploid, if the ISM holds at least approximately, and if the sequencing reads are obtained without bias from each homologous chromosome, then for each variant site, reads are sampled with equal coverage (call it *r/*2) from the chromosome with a variant and from the other without a variant. If the reads covering this site are randomly sampled from *n* cells, with the variant present in *k* cells, then the number *Z* of variant reads will be distributed as *Z*|*r* ~ Binomial 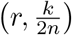. If the ploidy is equal to *P*, then *Z*|*r* ~ Binomial 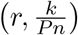. Therefore, (14) will stay the same, except for a change in the parameter of the binomial. It may be even modified to accommodate irregular ploidy (or copy number variation) along the chromosomes, although this will make the expressions more complicated.

It is very interesting to consider how the expectation of Ω(*x*_1_, *x*_2_), the number of mutations with sampling frequencies within (*x*_1_, *x*_2_] will be affected by change of ploidy *π*, if the expected SFS has the form as in Equ. (12). It is not difficult to conclude that the binomial humps will be transformed by *p*_*σ*_ → *p*_*σ*_/*P*.

It is less intuitive what the effect will be on the neutral (GT) part of the spectrum. As seen in the Figure (12), it will be simply multiplied by 1/*P*. This is likely related to the fact that under the ISM, mutated reads are sampled from the “1/*P* fraction” of the genome, so the effect is as if the mutation rate were multiplied by 1/*P*. This has to be reflected by a purely mathematical property of the GT spectrum, which is computationally evident, but not yet demonstrated rigorously. To make the property more specific: if the expected GT SFS is denoted *q*_*nk*_, *k* = 1 …, *n* − 1, then

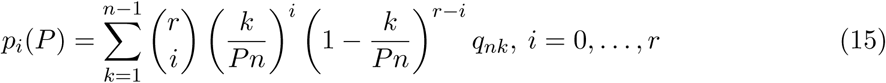

is inversely proportional to *P*.

### Driver mutations

In some cases, specific driver mutations identified as common in a given type of cancer can be found in a hump of the SFS. This is illustrated in the lung cancer cases TCGA-86-A4D0 and TCGA-62-A46O depicted in Figures 8 and 9. These results will be more comprehensively explored in another publication.

### Biologically meaningful parameters

As mentioned earlier, we can estimate a small number of aggregate parameters, which are functions of growth and mutation rates and the size of the background haplotypes of the emerging clones, as well as the proportions of these clones in cell population. The difficulty with interpretation of these parameters is illustrated best by the example of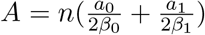. Suppose that we assume that mutation rate in the emerging clone 1 is the same as in clone 0. We still have to consider differences in growth rates between the two clones. These can be related to the proportion *p*_1_ of clone 1 (which is estimable), but the ages of the clones would have to be assumed.

### Dissection of humps and tumor evolutionary history

We return to the question discussed in Williams et al. (2018), namely the truncal mutations. If we assume there are *K*_0_ of these, we obtain the following augmented version of (12):

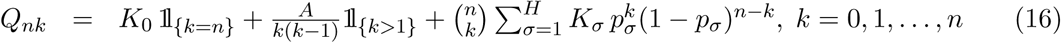

After transformation accounting for sampling and ploidy (as in Equ. (15), with *q*_*nk*_ replaced by *Q*_*nk*_), we see that the center GT term becomes one of the left-skewed profiles in Figure 12, the truncal term becomes a *K*_0_ Binomial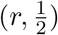 hump, and the right hand-side humps are transformed but retain their original masses *K_σ_, σ* = 1, *…, H*. In most cases analysed by us (see Figs. 6, 7, 8, and 9), we notice that all estimated hump masses are comparable to each other. If one of them corresponds to the truncal hump, this means that the ancestral cell of the tumor already acquired a very large mutation count. If the mass of this hump is approximately equal to 50% of all mutations, then this assertion might be consistent with the hypothesis of Tomasetti and Vogelstein (2015), who estimate that as many as 50% of mutations arise before transition to malignancy. It cannot be generally excluded that all the humps are truncal, corresponding to mutations in regions with different ploidies. However, for any regular ploidies *P*, the humps can be centered at (2*P*)^*−*1^ or to the left of this value contamination by reads from normal tissue is a problem. Probably only serial (in time) genome sequencing will allow to distinguish between this possibility and our model of secondary clones.

**Figure 12:**
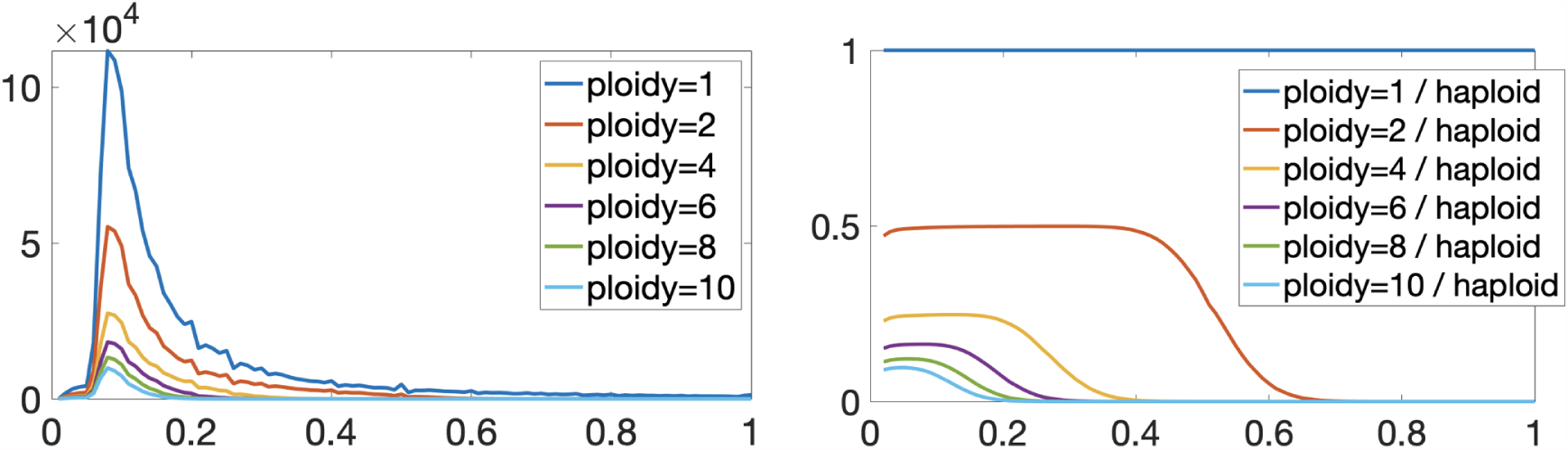
Transformation of sampled neutral GT spectrum under ploidy change

### Missing mutations

Based on (16), the total mass of the SMS is to good accuracy equal to

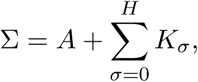

which is clear if we notice that 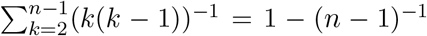. Table 3 indicates that this is many times more that the total number of mutations found in the sample. This result is understandable, if one considers that the fit producing parameter estimates were obtained using data pre-processing that makes the estimates sensitive only to the terms of of the GT-spectrum with *k ≥ L*. We may accept Σ as a crude estimate of the total count of point mutations in the tumor sample employed for sequencing.

### Single-cell sequencing data

The spread of new technologies will lead to breakthroughs in understanding of mutations and other genome transformations in cancer cells. Currently, we are witnessing a rapid expansion of single-cell DNA sequencing methods, such as described in Zahn et al. (2017).

With relatively low coverage, VAF values can be estimated reliably since they only may assume values from a spectrum *k/P*, where *k* = 1, *…, P*, if CNV or local ploidy at the given site is equal to *P*. This also allows us to infer in principle whether a substitution event at a given site preceded a chromosomal rearrangement or the other way around. However, it is unlikely that single-cell sequencing of a single snapshot of the tumor alone will bring a better understanding of evolutionary dynamics of cancer cell populations. This requires taking serial samples of DNA, which is still difficult at large scale.

### Recurrent mutations

The hypothesis underlying the methods in this paper is that mutations arise only once, so that recurrent mutations at any site are impossible. Whether this is satisfied or not depends on the mutation rate predominant at a given region of the genome. In the context of autosomal genomes, Kuipers et al. (2017) showed, using single-cell data, that it is highly unlikely that cancer cells do not feature recurrent mutations. Based on this possibility, Cheek and Antal (2018) provide a theory of SFS spectra that includes the possibility of recurrent mutation.

## A Appendix

### A.1 Simulating the Moran model with exponentially varying population size

Following Griffiths and Tavaré (1998), define 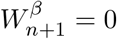, and for *j* = *n, n* − 1, …, 2, let

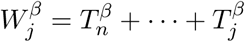

be the time until the sample from the exponentially growing population has *j* − 1 distinct ancestors. Then the waiting times 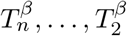 have distributions such that the conditional distribution of 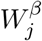, given 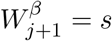 has density function

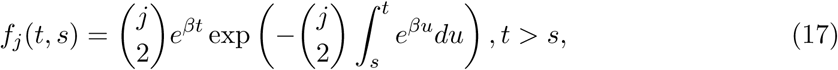

It follows from (2) that 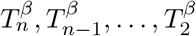 may be simulated via

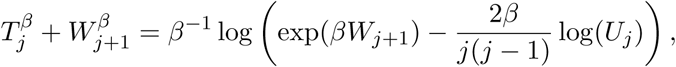

where *W*_*j*_ = *T*_*n*_ + … + *T*_*j*_, the *T*_*j*_ are independent exponential random variables with 𝔼*T*_*j*_ = 2/*j*(*j* − 1), and the *U*_*j*_ are independent, identically distributed (iid) uniform random variables on (0,1).

### A.2 Simulating the SFS of the lbdp

We follow the coalescent point process description given in Popovic (2004) and Lambert (2010). The individuals in the sample, numbered from 0 to *n −* 1, constitute tips of a rooted binary tree (with root at the bottom) ordered so that descendant nodes of a given node are placed on the right of that node, but on the left of the descendant nodes of any node that precedes it (in the sense that it is closer to the root of the tree). We limit ourselves here to the case of the coalescent process for the lbdp started at time *x* before present. Lambert (2010) and Lambert and Stadler (2013) have made this methodology much more general, but this is not crucial for the application we consider.

We define random variables *H*_0_, *H*_1_, *…* as the consecutive coalescence times. Following Lambert (2010), Theorem 5.4, they form a sequence of iid random variables with tail *W* (*t*)^*−*1^, killed at the first value larger than *x*. Hence, conditional on having *n* tips, they form a sequence of *n* iid random variables with tail *W* (*t*)^*−*1^ conditioned on being smaller than *x*.

We proceed to develop a simulation algorithm in the following steps (see Figure 13 for an example):

1. Let {*H*_*l*_}_*l*=0,…,*n*−1_ be the *H*_*i*_ ordered from largest (*H*_0_ = *x*) to smallest (*H*_*n*−1_). Let {*P*_*l*_}_*l*=1,…,*n*_ be the tip corresponding to the *H*_*l*_. Therefore *P*_*l*_ = *i* means that the tip *i* has the (*l* + 1)-th largest *H*_*i*_.
2. The times between ranked coalescence events (counting from the present in reverse time) are *H_n−_*_1_, *H_n−_*_2_ *− H_n−_*_1_, *…, H*_0_ *− H*_1_.
3. Let {*K*_*ij*_}_*i,j*=0,…,*n*−1_ denote the count of tips of the tree subtended by the *j*-th branch at the *i*-th level of the tree. The matrix *{K_ij_}* corresponding to the tree of Figure 13 is depicted in Figure 14.

**Figure 13:**
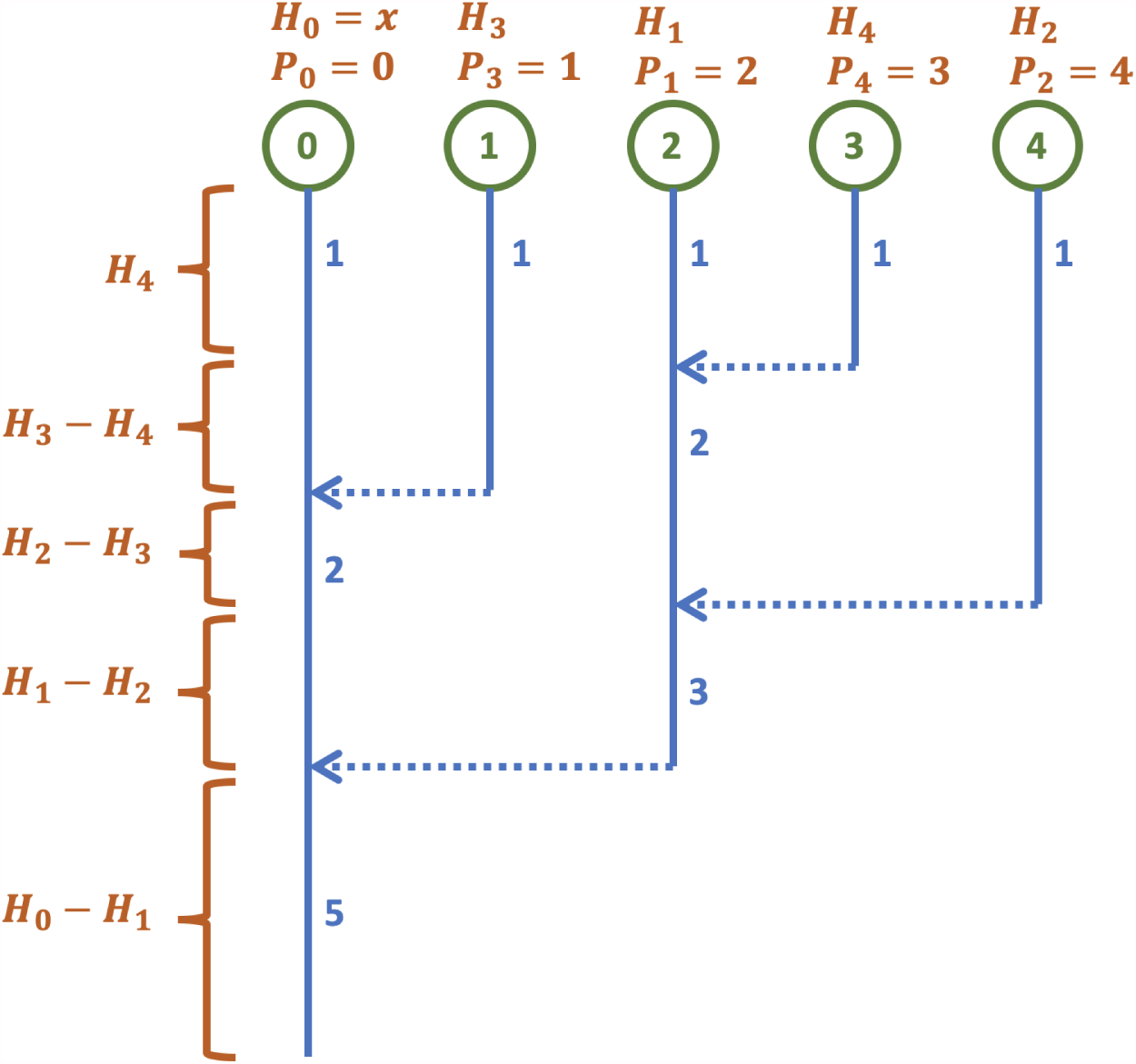
Schematic of a coalescent tree for the ldbp

The following algorithm recovers the terms of matrix {*K*_*ij*_}

1. *i* = 0: *K*_0*j*_ = 1, *j* = 0, *…, n −* 1
2. *i* = 1 *→ n −* 1:
  a. *l* = *n* − *i*
  b. *k* = max{*j*: *K*_*i*−1,*j*_ ≠ 0, 0 ≤ *j* < *P*_*l*_}
  c. 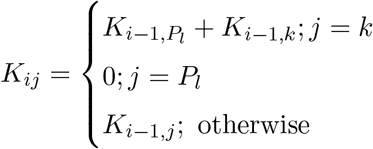

**Figure 14:**
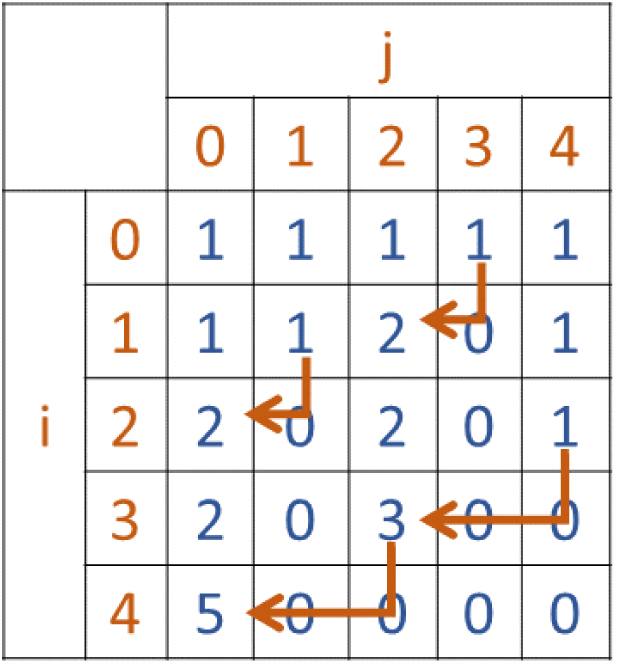
Matrix {*K*_*ij*_} corresponding to the tree of Figure 13

Finally, if the mutation process is Poisson with intensity *θ*, then a single realization of the SFS is generated as

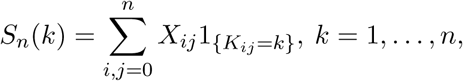

where the independent rv’s *X*_*ij*_ have Poisson distributions with parameters as follows

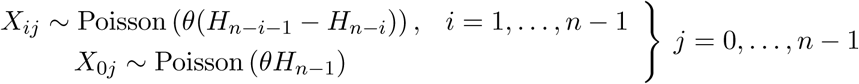

### A.3 Exact results for the SFS for the lbdp model

An explicit form of (7) is obtained by elementary derivation if *r >* 0. It boils down to computing the two definite integrals

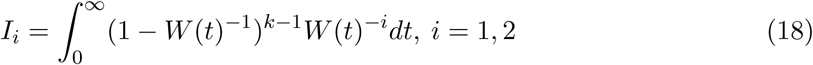

Substituting *z* = 1 *− W* (*t*)^*−*1^ = (*W* (*t*) *−* 1)*/W* (*t*), with inverse *W* (*t*) = (1 *− z*)^*−*1^ and *dt* = *r*^−1^ (1 *− z*)^−2^((1 *− z*)^−1^ − *α*)^−1^ d*z*, and noting that 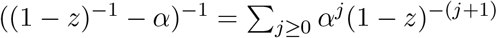, which converges under our assumptions, we obtain that

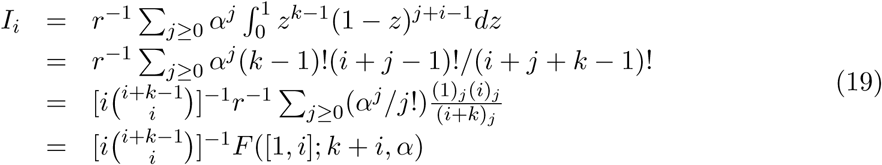

where 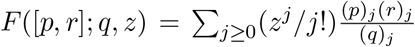 belongs to the hypergeometric family of functions (Abramowitz and Stegun, 1964) and

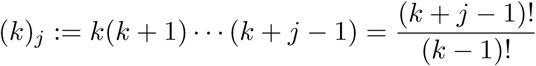

Combining these expressions with Equ. (7) we obtain

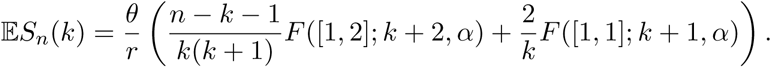

For the special case of pure birth process (*d* = 0) sampled in entirety (*p* = 1), which implies *α* = 0, we obtain

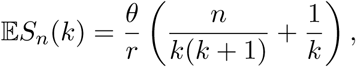

which is consistent with direct computation under *W* (*t*) = *e*^*rt*^. This expression suggests that for large *n*, Lambert SFS behaves as GT SFS for small *k*, i.e. it decays as *k*^−2^, but then as *k* approaches *n −* 1, it decays approximately as *k*^−1^.

It is interesting that the hypergeometric *F* ([1, 2]; *i, α*) can be expressed in the terms of finite sums of elementary functions for *α* ∈ (0, 1) and integer *i* ≥ 2. In particular *F* ([1, 2]; 2, *α*) = *α*(1 *− α*), and the following integral representation is valid for *i* ≥ 3

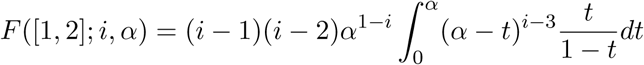

Based on this, *F* ([1, 2]; 3, *α*) = *−*2*α^−^*^2^(ln(1 *− α*) + *α*), and for larger *i* we obtain finite sums of products of polynomials and logarithmic terms.

